# Connectivity and network state-dependent recruitment of long-range VIP-GABAergic neurons in the mouse hippocampus

**DOI:** 10.1101/364547

**Authors:** Ruggiero Francavilla, Vincent Villette, Xiao Luo, Simon Chamberland, Einer Muñoz-Pino, Olivier Camiré, Kristina Wagner, Viktor Kis, Peter Somogyi, Lisa Topolnik

## Abstract

GABAergic interneurons in the hippocampus provide for local and long-distance coordination of neurons in functionally connected areas. Vasoactive intestinal peptide-expressing (VIP+) interneurons occupy a distinct niche in circuitry as many of them specialize in innervating GABAergic cells, thus providing network disinhibition. In the CA1 hippocampus, VIP+ interneuron-selective cells target local interneurons. Here, we discovered a novel type of VIP+ neuron whose axon innervates CA1 and also projects to the subiculum (VIP-LRPs). VIP-LRPs showed specific molecular properties and targeted interneurons within the CA1 area but both interneurons and pyramidal cells within subiculum. They were interconnected through gap junctions but demonstrated sparse spike coupling in vitro. In awake mice, VIP-LRPs decreased their activity during theta-run epochs and were more active during quiet wakefulness but not coupled to sharp-wave ripples. Together, the data provide new evidence for VIP interneuron molecular diversity and functional specialization in controlling cell ensembles along the hippocampo-subicular axis.

## Introduction

Understanding brain computations during different cognitive states requires identifying cell types, their connectivity motifs and the recruitment patterns under different behavioural conditions. GABAergic inhibitory neurons play a pivotal role in cortical computations through gain control, sensory tuning and oscillatory binding of cell ensembles (Atallah and Scanziani, 2009; Lapray et al., 2012; Lovett-Barron et al., 2012; Royer et al., 2012). However, understanding cortical inhibition has been a challenging task as this process is executed through a diverse group of local and long-range projecting (LRP) GABAergic neurons (Soltesz, 2006). Many types of GABAergic cells that have been identified by earlier investigations remain functionally uncharacterized. This is especially the case for sparse cell types, which represent a minority of cortical neuronal population and, therefore, have not been frequently sampled in blind electrophysiological recordings. In particular, until recently, very little has been known about the functional organization of GABAergic cell types that are specialized in the selective coordination of inhibitory interneurons. These so-called interneuron-selective (IS) cells express vasoactive intestinal peptide (VIP) alone or in combination with calretinin (Acsády et al., 1996a, b). They originate from the caudal ganglionic eminence and are the last cells to integrate into the cortical habitat (De Marco Garcia et al., 2011; Miyoshi et al., 2015), where they innervate many different types of local interneurons, including the somatostatin (SOM+), calbindin (CB+), parvalbumin (PV+), VIP (VIP+) and calretinin (CR+) expressing GABAergic cells (Acsády et al., 1996a, b; David et al., 2007; Staiger et al., 2004). Development of novel transgenic and optogenetic technologies allowed to investigate how these cells can coordinate the operation of cortical microcircuits (Ayzenshtat et al., 2016; Fu et al., 2014; Jackson et al., 2016; Lee et al., 2013; Pfeffer et al., 2013; Pi et al., 2013). A common finding between different cortical regions is that VIP+ IS cells suppress some local interneuron activity during complex behaviors, including visual processing (Ayzenshtat et al., 2016; Jackson et al., 2016; Pfeffer et al., 2013), locomotion (Fu et al., 2014), and reward-associated learning (Pi et al., 2013), thus leading to network disinhibition. However, similar to other GABAergic cells, VIP+ neurons are diverse in properties (Acsády et al., 1996a, b; Bayraktar et al., 2000; Porter et al., 1998; He et al., 2016) and, likely, in circuit function. Yet, no attempt has been made for a detailed physiological and functional analysis of morphologically defined subtypes of VIP+ interneurons. The hippocampal CA1 inhibitory circuitry can be considered one of the best characterized so far. Indeed, over the last three decades, the findings of multiple laboratories have culminated in a detailed wiring diagram of hippocampal CA1 GABAergic circuitry, with at least 21 inhibitory cell types identified to date (for review see Somogyi, 2010). Hippocampal CA1 VIP+ interneurons constitute two functionally different GABAergic cell populations: basket cells (BCs; Somogyi et al., 2004) and IS interneurons (IS2 and IS3 cells; Acsády et al., 1996a), which can modulate the activity of principal cells (PCs) or of different types of CA1 interneurons with a different degree of preference (Karson et al., 2009; Tyan et al., 2014). VIP+ BCs (VIP-BCs) can co-express cholecystokinin (CCK) and, in addition to targeting PC somata, can contact PV-positive BCs, indicating that VIP-BCs can exert both inhibitory and disinhibitory network influences (Karson et al., 2009). In contrast, the VIP+ IS interneurons prefer to contact inhibitory interneurons (Acsády et al., 1996a), and modulate interneuron firing properties (Tyan et al., 2014). Although disinhibition can be a common mechanism of hippocampal computations necessary for the induction of synaptic plasticity and memory trace formation and consolidation (for review see Letzkus et al., 2015), current findings indicate that its effect is mostly local due to the local innervation of hippocampal inhibitory microcircuits through VIP+ interneurons (Tyan et al., 2014). Interestingly, anatomical data point to the existence of long-range circuit elements that could account for cross-regional disinhibition between the hippocampus and functionally connected areas: CA1 SOM-or muscarinic receptor 2 (M2R)-expressing GABAergic cells innervate hippocampal inhibitory interneurons and can project to several cortical and sub-cortical areas, including the rhinal and retrosplenial cortices, subiculum (SUB) and medial septum (MS) (Gulyás et al., 2003; Jinno et al., 2007; Miyashita and Rockland, 2007; Fuentealba et al., 2008; Melzer et al., 2012). Despite the considerable recent interest in LRP GABAergic neurons, very little is currently known about the connectivity and function of these cells during different network states in awake animals. Here, we reveal a novel subtype of VIP-expressing LRP (VIP-LRP) GABAergic neuron that exhibits a specific molecular profile and innervates, in addition to the hippocampal CA1, the SUB, with region-specific target preference. Functionally, VIP-LRP cells correspond to theta-off cells (Buzsáki et al., 1983; Colom and Bland, 1987) as they decrease their activity during theta-run epochs associated with locomotion and exhibit high activity during quiet wakefulness. The identification of this circuit element reveals a new mechanism for the behaviour- and network-state-dependent inter-regional coordination of activity within the hippocampal formation.

## Results

### Long-range-projecting VIP-expressing neuron in the CA1 hippocampus

To characterize the electrophysiological and morphological properties of VIP+ interneurons in the hippocampal CA1 area, we first performed patch-clamp recordings from VIP+ cells in acute slices obtained from VIP-eGFP mice (Figures 1A, S1, S2G, S2I; see also Tyan et al., 2014 for characterization of the GFP expression in this mouse strain). Following biocytin labelling, 97 VIP-GFP+ interneurons were visualized and identified as BCs, IS3 cells or as novel LRP neurons (VIP-LRP; Figures 1B, S1B; Table S1). The VIP-LRP neurons typically occurred at the oriens/alveus (O/A) border and had horizontal sparsely spiny dendrites, which were mostly restricted to the stratum oriens (Figure 1B). Their axon formed a local arbor in O/A and extended slightly into strata pyramidale (PYR) and RAD of the CA1 region. The main axon was partially myelinated and travelled outside the CA1, giving rise to a large axon cloud in the proximal SUB (Figure 1B; n = 40 cells out of 78 biocytin-filled O/A VIP-GFP+ interneurons). Thus, in contrast to VIP-BCs and to IS3 cells, the axon of VIP-LRP neurons occupied two major areas: CA1 O/A and SUB (Figure 1B; 1E). The total length of the axonal arbor in a 300-µm slice was between 7,704 and 36,167 µm (no shrinkage correction). The axon length occupying the CA1 vs SUB was 2,000–20,000 (median ± SD: 10,184 ± 5,002) and 1,000–26,000 (median ± SD: 5,041 ± 6,740) µm, respectively; the large variability was likely due to a different degree of the axon preservation in slices (Figure 1E; n = 10 cells). These cells showed a regularly spiking firing pattern and a membrane potential ‘sag’ in response to a hyperpolarizing step to –100 mV (Figures 1C, S1A). Furthermore, their intrinsic membrane properties were similar to those of VIP-BCs [except for the sag and the fast afterhyperpolarization (AHP) amplitude] but differed in many parameters from IS3 cells (Table S1). To provide additional evidence for the presence of SUB-projecting VIP+ neurons in the CA1 O/A, we injected red RetroBeads in the SUB of VIP-eGFP mice. In addition to pyramidal cells, bead-labelled VIP-GFP-positive neurons were detected in the CA1 O/A area (Figure 1D, left), thus confirming that a population of O/A VIP+ neurons sends long-range axons to the SUB. As to their molecular profile, all VIP-LRP cells tested with an axon reaching SUB were immunopositive for muscarinic receptor 2 (M2R; n = 7/7 cells; Figure 1D, right), thus identifying the M2R as an additional molecular marker of VIP-LRPs. Furthermore, a large fraction of M2R+/VIP-GFP+ cells co-expressed CB (29/53 cells; Figure S2A, S2I) but were negative for CCK (Figure S2D, S2I), nitric oxide synthase (NOS; Figure S2E, S2I), CR (Figure S2F) and SOM (Figure S2H, S2I). This was in contrast to VIP-BCs and IS3 cells, which co-expressed CCK or CR, respectively, and were negative for M2R (Figure S2C). Overall, 50% of VIP-GFP+ cells in the CA1 O/A of VIP-eGFP mouse were co-expressing M2R (Figure S2I), corresponding to the VIP-LRP population. In the rat, trilaminar cells projecting to the subiculum are rich in M2R in the somato-dendritic membrane and are innervated by presynaptic mGluR8-positive terminals (Ferraguti et al., 2005). We tested if the VIP-GFP-M2R-positive cells in the mouse received mGluR8+ input and found that most VIP-GFP-M2R-positive cells in O/A were decorated by mGluR8+ terminals, some of which were themselves VIP-positive as in the rat (Figure 1F). Some M2R+ neurons in O/A were not immunoreactive for VIP and GFP, and not all VIP-GFP+ neurons showed M2R immunoreactivity (Figure S2I), pointing to additional molecular diversity within M2R+ and VIP+ neuronal populations.

**Fig. 1.**
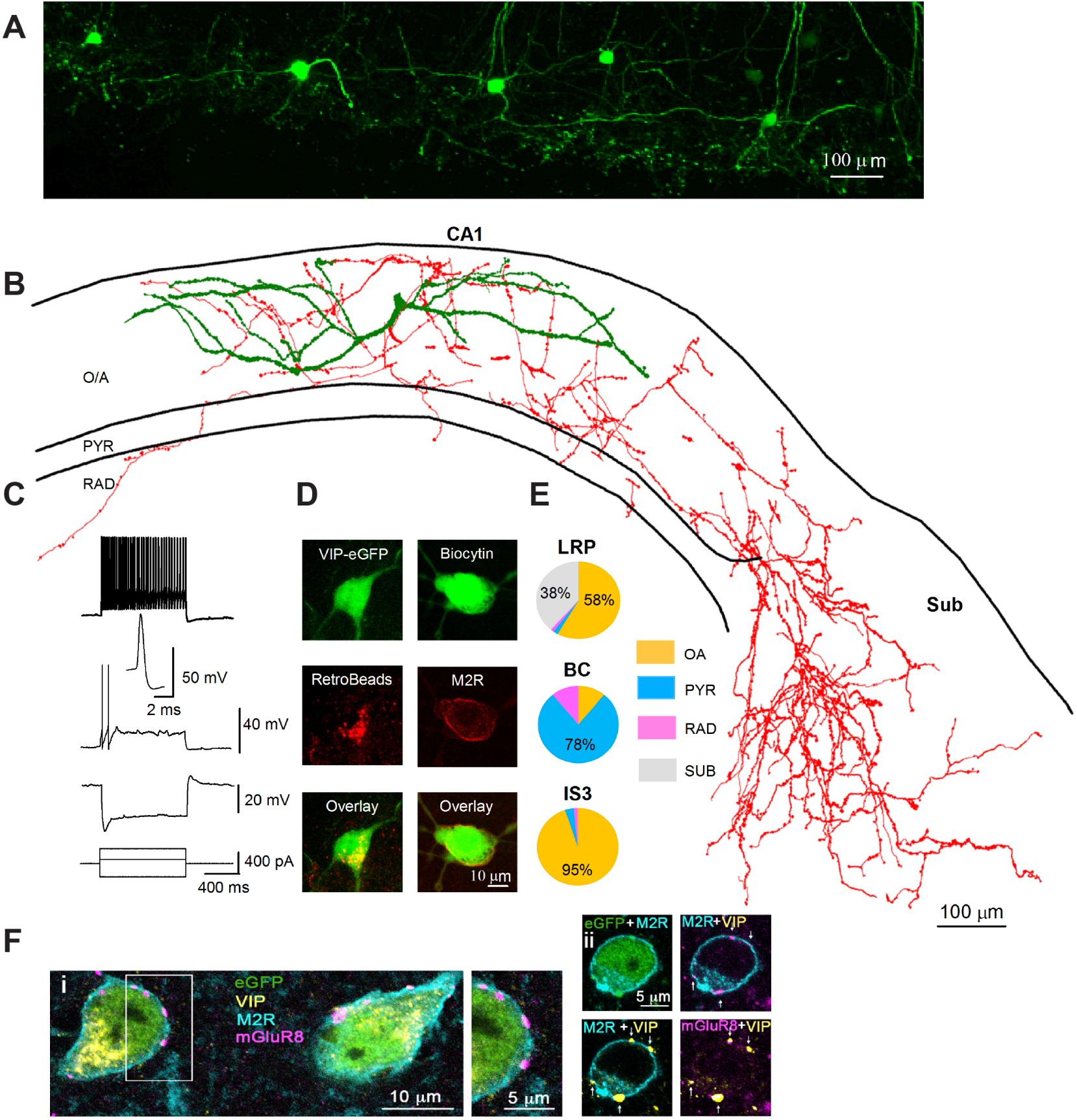
Identification of VIP-LRPs in the VIP-eGFP mouse. See also Figure S1, S2 and Table S1, S2. (A) Two-photon image (maximal projection of a z-stack of 200 µm height) of the CA1 area from an acute hippocampal slice (300 µm) of a VIP-eGFP mouse showing the location of GFP cell bodies, axons and dendrites in the O/A area of CA1. (B) Reconstruction (the axon is shown in red, the dendrites are shown in green) of a VIP-LRP cell that was recorded and filled with biocytin in a slice obtained from a VIP-eGFP mouse. (C) Representative voltage responses of a VIP-LRP to hyperpolarizing (−240 pA), and depolarizing (+80 pA and +280 pA) current injections, with an inset illustrating the first spike evoked by +80-pA current pulse. (D) Confocal images showing RetroBeads labelling of a VIP-LRP soma (left) after injection in subiculum and immunoreactivity for M2R in a VIP-positive neuron labelled with biocytin (single focal plane, right). (E) Pie charts illustrating the mean axonal distribution in different layers (based on axon length obtained following reconstruction in Neurolucida) for groups of cells corresponding to 3 different cell types: VIP-LRP (n = 10), VIP-BC (n = 5) and IS3 cell (n = 6). OA, oriens-alveus; PYR, stratum pyramidale; RAD, stratum radiatum and SUB, subiculum. No axon was detected within stratum lacunosum moleculare (LM) for the three cell types. Statistically significant differences in the axon distribution between VIP-LRP and VIP-BCs, VIP-LRP and IS3, and VIP-BCs and IS3 at **p< 0.01, one-way ANOVA followed by Tukey’s test. (F) VIP-LRP cells, identified by somato-dendritic membrane M2R immunoreactivity (blue), are innervated by terminals rich in presynaptic mGluR8 (purple). Single optical slices (0.45 mm thick) of confocal images of quadruple immunoreactions as indicated: i, right, framed area at higher magnification; ii, four VIP+ terminals (arrows) show mGluR8 immunoreactivity.

### Local connectivity of VIP-LRP cells

To determine the VIP-LRP physiological function, we next examined its local connectivity using simultaneous paired recordings and electron microscopic (EM) analysis (Figure 2). Dual whole-cell patch clamp recordings showed that out of 118 attempts, 33 pairs of VIP-LRPs and CA1 O/A interneurons were connected synaptically, and no connection was found with CA1 PCs (Figure 2I, see also Figure 4H and Figure 7J for additional connectivity studies). Among the VIP-LRP targets, we identified different types of dendritic inhibitory cells (DT-INs), such as O-LM (Figure 2A–2E) and bistratified cells (BIS; Figure 2F), and the perisomatic terminating interneurons (ST-INs), such as BCs (Figure 2G). As O-LM cells were the most frequent target, we characterized the VIP-LRP to O-LM cell connection in more details (Figure 2A–2E). The VIP-LRP synapses occurred on dendritic shafts of OLMs (distance from soma: 63.4 ±19.8 µm, n = 7 pairs; Figure 2A), had mean unitary inhibitory postsynaptic current (uIPSC) amplitude of 16.3 ± 2.4 pA (0 mV holding potential) and a failure rate of 60.1 ± 4.1 % (n = 8 pairs). uIPSCs had slow kinetics consistent with dendritic location of synapses (Fig. 2D). During repetitive VIP-LRP firing, uIPSC showed no change at 10 to 50 Hz but summated efficiently at 100 Hz (Figure 2C, 2E). Apart from the uIPSC amplitude, which was higher in BISs (32.1 ± 6.2 pA, n = 5 pairs; Figure 2J), the properties of VIP-LRP synapses were similar among different postsynaptic targets (Table S3), and, also, did not differ significantly from synapses made by IS3 cells on O/A interneurons (Tyan et al., 2014).

**Fig. 2.**
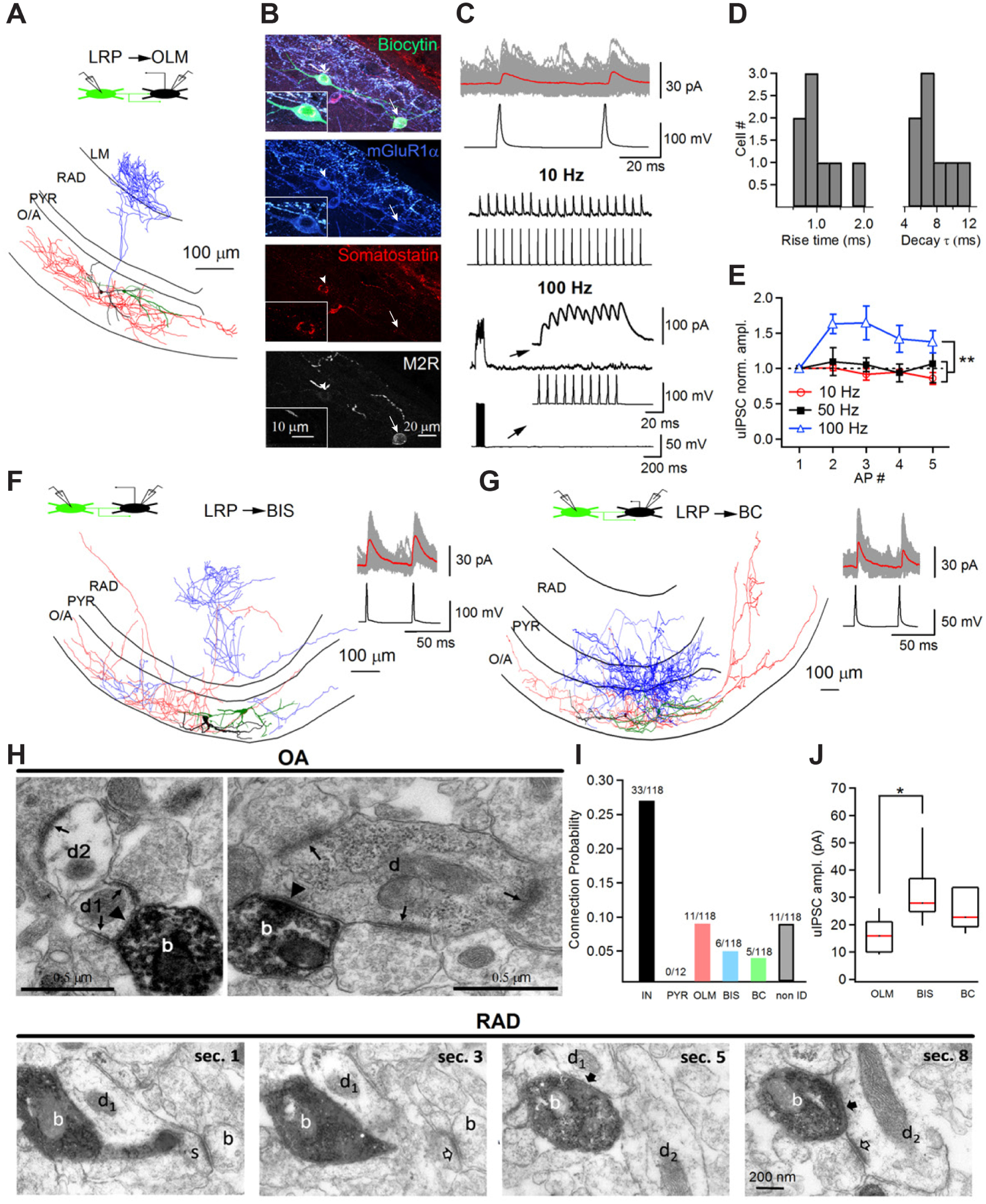
VIP-LRPs provide inhibition to different subtypes of CA1 O/A interneurons. See also Table S3. (A) Reconstruction of a synaptically connected pair of VIP-LRP and oriens-lacunosum moleculare (O-LM) interneurons. The axon of the presynaptic VIP-LRP cell is shown in red and its dendrites are shown in green. The axon of the postsynaptic O/A interneuron is shown in blue, with its dendrites shown in black. The inset shown on top illustrates schematically the configuration of the recording. (B) Post hoc immunohistochemical analysis of a different synaptically connected pair of VIP-LRP (immunoreactive for M2R, arrow) and O-LM cell (arrowhead) with insets showing immunoreactivity of postsynaptic O-LM cell. (C) Representative traces of voltage-clamp recordings of uIPSCs in an O-LM cell at 0 mV (100 consecutive traces are shown in gray and the average is shown in red, top) evoked by two APs through a current injection in a presynaptic VIP-LRP cell (top), and examples illustrating the changes in uIPSC amplitude during different frequencies of firing of VIP-LRPs: 10 Hz (middle) and 100 Hz (bottom). Insets at the bottom show an expanded view of the uIPSCs during firing of VIP-LRP at 100 Hz. (D) Cumulative histograms of uIPSC rise time and decay time constant in O-LM cells (n = 11 pairs). (E) Summary plot showing changes in uIPSC amplitude in O-LM cells during different frequencies of firing of VIP-LRPs. Statistically significant differences in the uIPSC amplitude between 100-Hz and 10-Hz presynaptic firing, and 100-Hz and 50-Hz presynaptic firing at **p< 0.01, one-way ANOVA followed by Tukey’s test. (F-G) Reconstructions of a synaptically connected pair of VIP-LRP and a putative bistratified cell (BIS; F) and of a VIP-LRP and basket cell (BC; G). The insets on the right show voltage-clamp recordings of uIPSCs at 0 mV in response to APs evoked by the current injection in the presynaptic VIP-LRP cell. (H) Top images show electron micrographs (EM) of two biocytin labelled boutons (b) of a GFP-positive horizontal interneuron recorded in stratum oriens. The cell had a main axon heading towards the subiculum in the white matter. The electron-opaque HRP reaction product marks the boutons, which form type-2 synapses (arrowheads) with a small (left d1) and a large diameter (right) dendritic shafts (d) of interneurons receiving mostly type-1 synapses (arrows) from unlabeled boutons in stratum oriens. Another interneuron dendritic shaft (d2) also receives a type-1 synapse. Bottom EM images correspond to four neighbouring sections (sec. 1, 3, 5 and 8) illustrating dendrites (d1, d2) in CA1 RAD as local postsynaptic targets of a VIP-LRP shown to project to the subiculum. The interneuron bouton (labelled with biocytin, white ‘b’) makes two type-2 synapses (solid arrows): with a spiny dendrite (d1, s), where the spine is receiving a type-1 synapse (open arrow), and with a dendrite (d2) receiving a type-1 synapse (open arrow) on the shaft. (I-J) Summary bar graphs illustrating the connection probability (I) defined as a ratio between the number of connected pairs of a given type to the total number of attempts and the uIPSC amplitude (J) for different postsynaptic targets.

To further validate the results of paired recordings, EM analysis of 39 synaptic junctions within the CA1 area made by 2 VIP-LRPs filled with biocytin was performed. The data showed that interneuron dendrites were frequent synaptic targets of VIP-LRPs. Of the 18 postsynaptic dendrites tested from the targets of one VIP-LRP, 16 originated from interneurons and 2 were unidentified (Figure 2H, top). The other VIP-LRP cell made synapses with 6 interneuron dendrites and 15 unidentified dendrites, some of which emitted spines (Figure 2H, bottom), suggesting that spiny interneurons or PCs could be among the VIP-LRP targets. Indeed, in optogenetic experiments (see below), we found that two out of 49 CA1 PCs examined received input from VIP-LRPs (Figure 7J, S7D). Taken together, these data indicate that, locally, VIP-LRPs prefer to target interneurons and constitute a novel type of IS cells in the hippocampus.

To investigate the possible connectivity within the VIP-LRP population, we performed dual whole-cell patch-clamp recordings from VIP-GFP+ pairs (Figure 3A). Our data showed that out of 36 attempts, 16 VIP-LRP pairs were connected through symmetric gap junctions with a coupling co-efficient of 0.11 ± 0.1 (Figure 3D, 3E), and one pair was connected synaptically. Electrotonic coupling between VIP-LRPs was blocked by the selective connexin-36 gap-junction blocker mefloquine (Cruikshank et al., 2004) (to 7.1 ± 3.8 % of control, n = 6 pairs; p < 0.01; paired t test; Figure 3D, 3E) or the broad-spectrum gap-junction blocker carbenoxolone (to 22.4 ± 14.4 % of control, n = 4 pairs; p < 0.01; paired t test; Figure 3E). Furthermore, the electrotonic signal conduction exhibited low-pass filter properties. As such, fast action potentials (APs) generated in cell 1 were strongly attenuated in cell 2, whereas the slow AHPs were better conducted, leading to a substantial hyperpolarization of cell 2 (Figure 3B; 3C; 3G, bottom right insets). We then examined how a sinusoidal excitatory input modulated at theta-like frequency can be integrated by the electrically coupled VIP-LRP neurons (Figure 3F–3H). We found that, when both cells were kept at rest, a sinusoidal input applied to cell 1 induced subthreshold synchronous fluctuations of the membrane potential of cell 2, but was not able to drive its firing (Figure 3F). When cell 2 was slightly depolarized to allow for spontaneous firing, the two VIP-LRPs could occasionally fire together but their coupling remained weak, and most spikes occurred asynchronously due to the AHP-associated inhibition of cell 2 (Figure 3G). The voltage fluctuation in cell 2 was significantly higher when spikes were not generated in cell 1 [voltage peak without AP in cell 1: 8.5 ± 0.1mV (inset black trace) vs voltage peak with AP in cell 1: 6.9 ± 0.1mV (inset red trace), n = 6 pairs; p < 0.01; paired t test; Figure 3G, bottom right inset]. When both cells received a synchronous theta-modulated excitatory input, their firing increased (to 112 % n = 6 pairs; Figure 3H), but the spike synchrony remained weak (Figure 3H, bottom). Together, these data indicate that electrotonic coupling between VIP-LRPs is unlikely to synchronize their recruitment in response to theta-like input. Whether this may be the case at a different firing frequency (Alvarez et al., 2002; Trenholm et al., 2014), activity-dependent state of the gap junctions or network size will need to be explored using computational modeling (Pernelle et al., 2018).

**Fig. 3.**
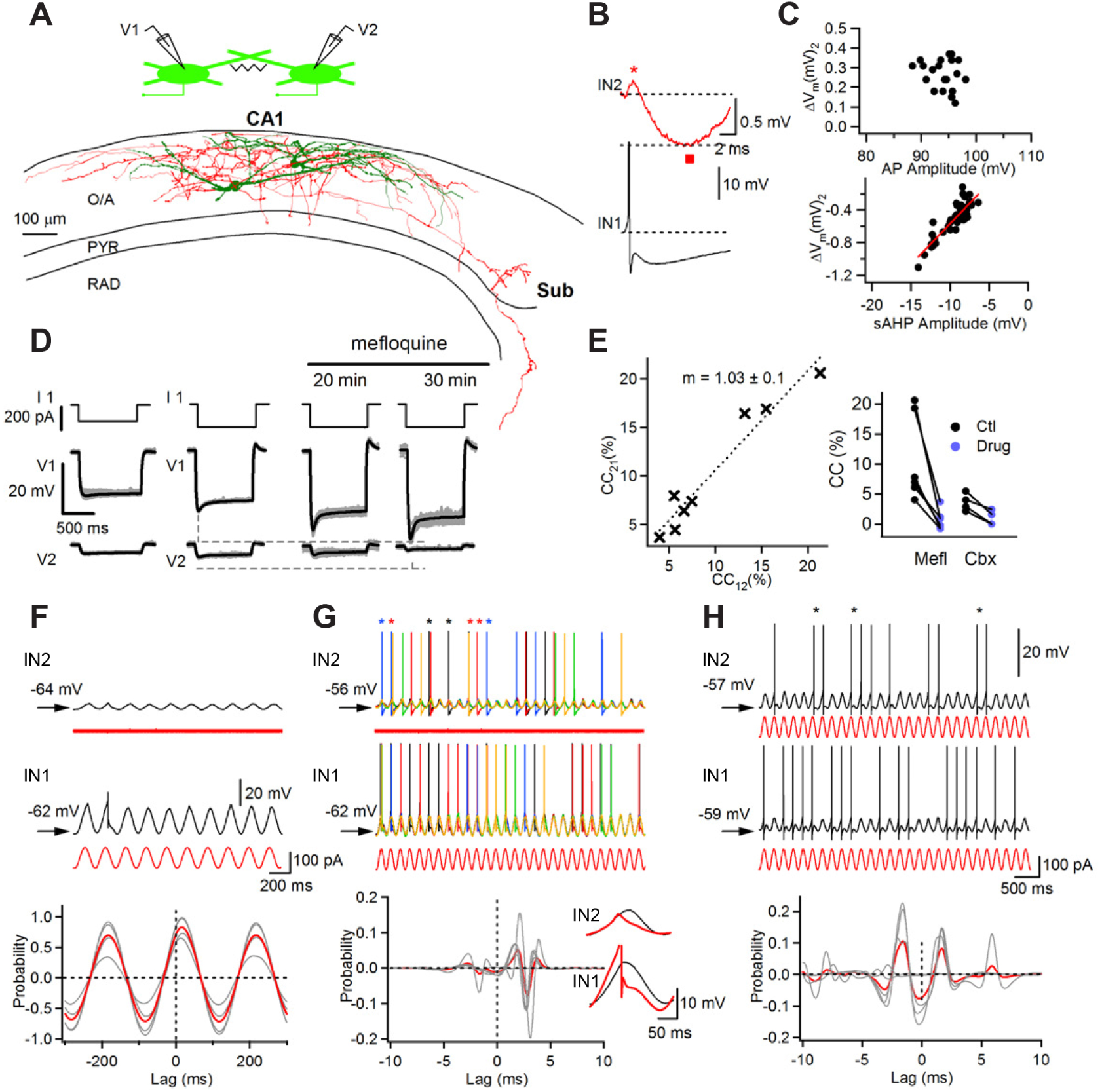
Electrical coupling between VIP-LRPs. (A) Schematic of the simultaneous recording of two VIP-LRPs and their reconstruction. (B) Representative example of the simultaneous recording of two VIP-LRPs with an AP initiated in the presynaptic cell (IN1, black trace) and a corresponding voltage response in the postsynaptic cell (IN2, red trace). The positive and negative components of the spikelet are shown with a star and square symbols, respectively. (C) Summary plots for a group of cells (n = 17) indicating changes in the postsynaptic Vm as a function of the AP amplitude (upper) or sAHP amplitude (bottom). Red line is a linear fit to the data points (r = 0.88, Pearson correlation) for a slow negative spikelet component associated with sAHP in the presynaptic cell. (D) Representative examples of voltage traces recorded in the VIP-LRP pair before and after the application of the gap-junction blocker mefloquine (100 µM). (E) Summary plots for the coupling coefficient (CC) exhibiting symmetry between different pairs (left; m, slope of the regression line ± SE), and for the gap-junction blockers’ effect (Mefl, Mefloquine; Cbx, Carbenoxolone) for a group of cells (right; Mefl: n = 6; Cbx: n = 4; p< 0.05; paired t test). (F) Voltage responses recorded in a VIP-LRP pair (at a subthreshold level for AP generation Vm) to a sinusoidal current (red trace, 5 Hz) applied to the presynaptic cell (IN1). Plot below shows cross-correlations in Vm fluctuations between the two cells for a group of pairs (n = 6), with red trace corresponding to the average data. (G) Voltage responses (five consecutive traces of different colors superimposed) recorded in the VIP-LRP pair with the postsynaptic cell (IN2) being depolarized to allow for spontaneous firing and a sinusoidal current (red trace) applied to the presynaptic cell (IN1). Stars of different colors above the IN2 traces indicate APs generated synchronously in two cells. Plot below shows cross-correlations in the AP occurrence for a group of pairs (n = 6), with red trace corresponding to the average data. Insets on the right show voltage responses in two cells with (red trace) and without (black trace) an AP generated in the presynaptic cell. Note a larger Vm fluctuation in the postsynaptic cell (IN2) in the absence of AP in the presynaptic cell (IN1). (H) Voltage responses recorded in a VIP-LRP pair to a sinusoidal current (red trace, 5 Hz) applied to both cells. Stars above the IN2 trace indicate APs generated synchronously in two cells. Plot below shows cross-correlations in the AP occurrence for a group of pairs (n = 6), with red trace corresponding to the average data.

### Distant connectivity of VIP-LRP cells

To determine the distant targets of VIP-LRPs in SUB, we conducted single-cell two-photon glutamate uncaging-based mapping of connections by combining the photoactivation of CA1 O/A VIPGFP+ cells and patch-clamp recordings of interneuron and PC targets (Figure 4E). As reported previously (Chamberland et al., 2010), two-photon uncaging (730 nm, 20–30 mW/180 ms laser pulses) of locally delivered MNI-Glu (micropressure pulses via a glass pipette positioned above the cell of interest; see Methods for details) triggered single spikes in VIPGFP+ interneurons, resulting in fast uncaging-evoked IPSCs (glu-IPSCs; delay onset: 4.8 ± 1.8 ms) in target cells in case of connection (Figure 4A–D). Depending on the number of VIP-GFP+ cells per slice, this approach allowed us the testing of several VIP-GFP+ connections to a given target (1 to 5; Figure 4A–D). First, consistent with our findings using paired recordings and ultrastructural analysis (Figure 2), in the CA1 area, VIP-LRPs were connected to O/A interneurons (n = 9 connections out of 31 tested/11 cells; Figure 4A, 4F, 4H). Glu-IPSCs were not detected in CA1 PCs (n = 0 connections out of 18 tested/10 cells; Figure 4B, 4F, 4H), confirming the interneuron preference of VIP-LRPs. Morphological analysis of postsynaptic interneurons filled with biocytin also confirmed that BIS and O-LM cells were among the preferential targets of VIP-LRPs within CA1 O/A (data not shown). In addition, post hoc immunohistochemical examination of connected neurons revealed that VIP-GFP+ cells that were connected to O/A interneurons were positive for M2R (Figure 4G), consistent with the neurochemical profile of VIP-LRPs (Figures 1D, 2B). Surprisingly however, in the SUB, VIP-LRPs innervated both interneurons (n = 8 connections out of 45 tested/26 cells; Figure 4C, 4F, 4H) and PCs (n = 11 connections out of 37 tested/24 cells; Figure 4D, 4F, 4H) identified based on their dendritic and axonal properties (see Figure 7H for reconstructed examples). These data highlight the region-specific target preference of VIP-LRPs. The glu-IPSC amplitude was similar between all CA1 and SUB targets (Figure 4H, bottom), although some target-specific differences were observed within a population of CA1 interneurons (Figures 2J, 7J; Table S3).

**Fig. 4.**
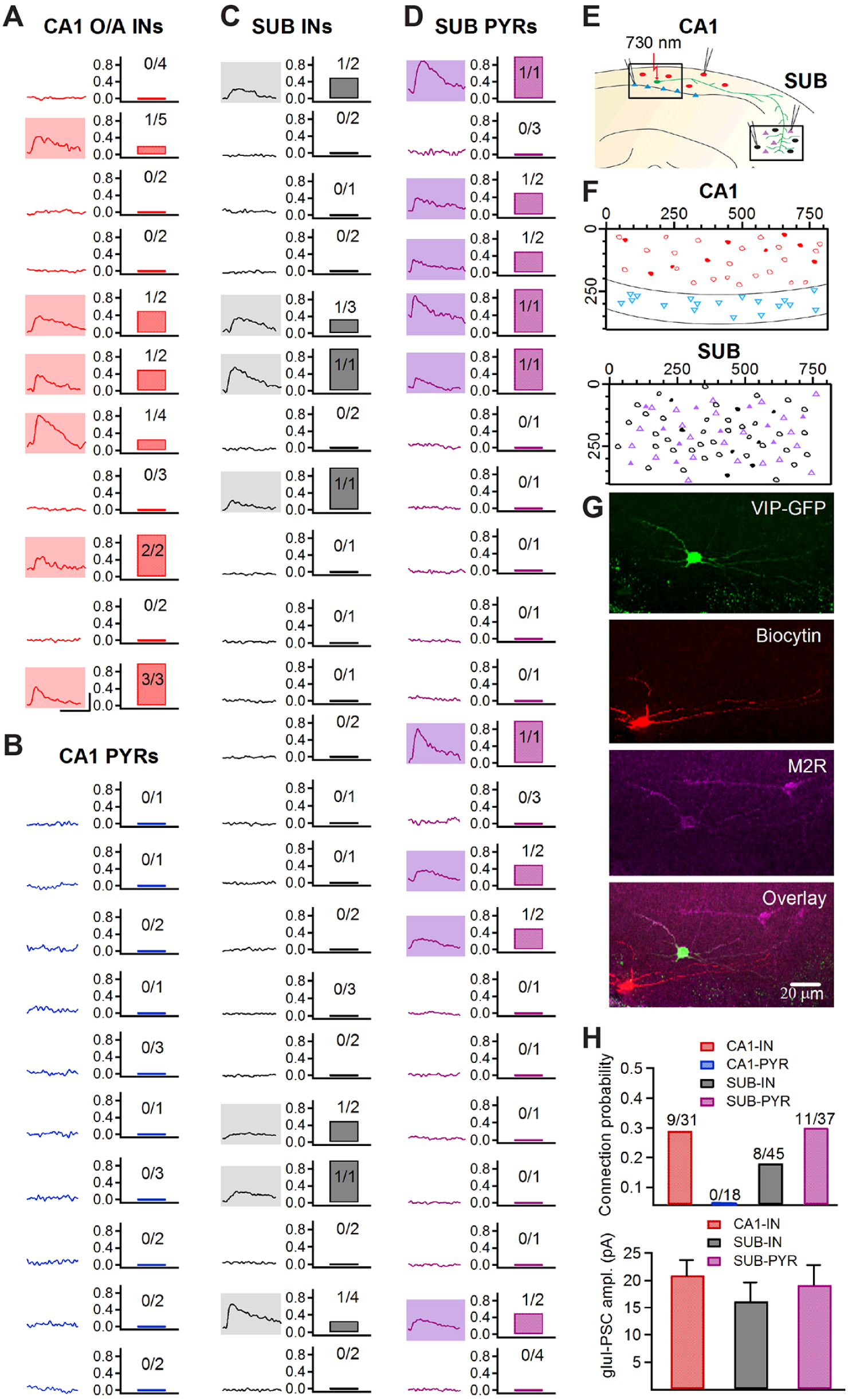
Two-photon glutamate uncaging-based mapping of local and distant axonal targets of VIP-LRPs. (A–D) Average traces of glu-IPSCs (Vhold: 0 mV) evoked by uncaging of MNI-Glu on VIP+ O/A interneuron somata (left) and the corresponding connection probability (right) in CA1 O/A interneurons (A), CA1 PCs (B), SUB interneurons (C) and SUB PCs (D). Each row corresponds to a single cell with the ratio of connections indicated at bar graphs. Each recorded cell was tested for receiving input from 1 to 5 VIP-GFP+ O/A interneurons. Traces with shadow area correspond to examples of glu-IPSCs: CA1 interneurons (n = 9 connections out of 31 tested/11 cells), CA1 PCs (n = 0 connections out of 18 tested/10 cells), SUB interneurons (n = 8 connections out of 45 tested/26 cells), SUB PCs (n = 11 connections out of 37 tested/24 cells). Scale bars (shown in A): 20 pA, 10 ms. (E) Schematic of simultaneous patch-clamp recordings from different CA1 and SUB targets and two-photon MNI-Glu uncaging on somata of VIP-GFP+ cells in CA1 O/A. (F) Summary spatial maps illustrating the density of connections within the CA1 (top) and SUB (bottom). Connected cells are shown as shaded symbols. Scales are in µm. (G) Post hoc immunohistochemical validation of connected interneurons confirmed that VIPGFP+ cells innervarting CA1 O/A interneurons were positive for M2R and, thus corresponded to VIP-LRPs. (H) Summary bar graphs showing the connection ratio (top) and the peak amplitude of glu-IPSCs (bottom) for different postsynaptic targets in CA1 and subiculum. The connection ratio is a ratio between the number of connections over the total number of tests for a given target type.

### Network-state-dependent recruitment of VIP-LRP cells in awake mice

To understand the functional role of VIPLRP cells, we performed in vivo two-photon calcium (Ca2+) imaging of VIP+ interneuron activity in parallel with local field potential (LFP) recordings from the contralateral CA1 hippocampus in head-restrained awake mice running on a treadmill (Villette et al., 2017). The Cre-dependent viral vector AAV1.Syn.Flex.GCaMP6f.WPRE.SV40 was delivered to the CA1 hippocampus of VIP-Cre mice to express Ca2+-sensitive protein GCaMP6f selectively in VIP+ neurons. The immunohistochemical analysis of VIP+ O/A neurons in VIP-Cre mice confirmed that VIP/M2R-co-expressing cells were present in the CA1 O/A (albeit at a significantly lower fraction when compared with VIP-eGFP mice: 7% in VIP-Cre vs 50% in VIP-GFP out of total VIP+ O/A cells; Figures S2, S3), likely due to the mouse strain differences (CD1 for VIP-eGFP vs C57BL/6J for VIP-Cre mice; Chia et al., 2005; Chen et al., 2006; Mekada et al., 2009). The M2R+ VIP O/A cells in VIP-Cre mice exhibited virus-driven GCaMP6f expression, showed a normal morphological appearance (Figure 5E) and were examined for the activity-dependent recruitment during different behavioural and network states.

**Fig. 5.**
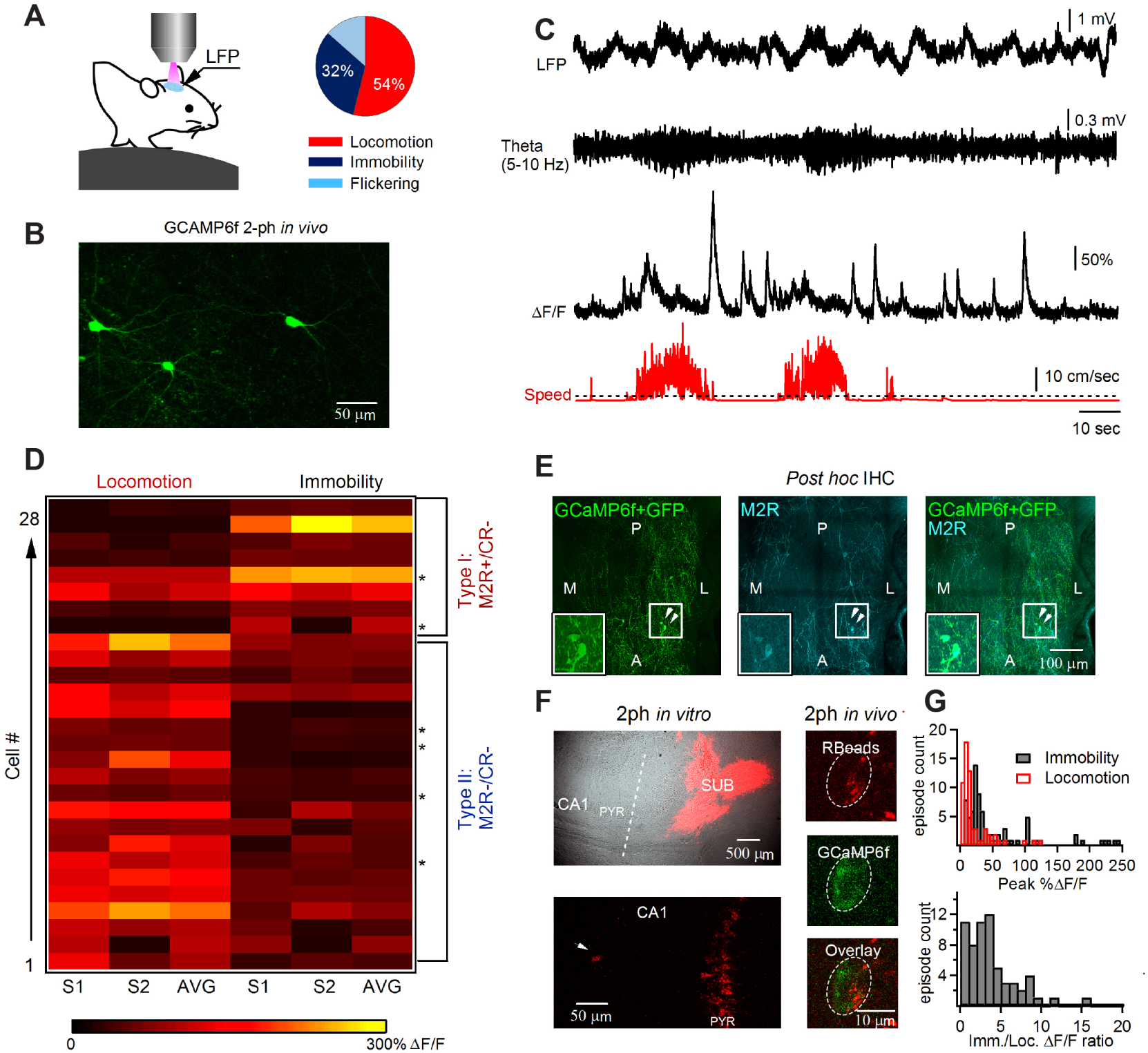
Imaging VIP-LRP activity in awake mice. See also Table S4. (A) Schematic of simultaneous two-photon Ca2+-imaging and LFP recordings in awake head-restrained mice (left) and a pie-chart illustrating the time distribution of different behavioural states (right). (B) Two-photon image of the GCaMP6f-expressing VIP cells in CA1 O/A (maximal projection of a 100-µm Z-stack) acquired at a high laser power to illustrate the cell morphology. (C) Representative traces of simultaneous LFP recording (upper raw trace and filtered for theta: 5–10 Hz) and somatic Ca2+-imaging from a VIP cell during different behavioural states (red trace; the higher animal speed during locomotion is reported as a high-frequency step pattern). (D) VIP O/A cell segregation based on the cell recruitment determined from the peak Ca2+-transient during locomotion and immobility. Two types of neurons were revealed: type I VIP cells (n = 8) were positive for M2R and negative for CR (2 out of 2 cells tested) whereas type II VIP-expressing cells (n = 20) were negative for both M2R and CR (4 out of 4 cells tested). S1 and S2 represent two independent imaging sessions and AVG is the average peak Ca2+-signal of the two sessions. (E) Confocal stitching of the hippocampal CA1 O/A imaging window used for two-photon image acquisition in vivo and processed for post hoc immunohistochemistry illustrating GFP (left) and M2R (middle) immunoreactivity as well as the overlay of the two (right). White arrows point to M2R-positive VIP cells targeted with GCaMP6f, which are illustrated as insets. Anatomical landmarks are indicated as following: A – anterior, P – posterior, M – medial, L – lateral. (F) Representative two-photon images obtained from hippocampal slices of VIP-Cre mice in vitro (left) and from the CA1 O/A VIP+ interneuron in vivo (right) illustrating the RetroBead injection site in SUB (left top, Dodt infrared scanning gradient contrast image superimposed with epifluorescence image for Retrobeads acquired with 4x lens), the retrogradely labeled CA1 PCs and an O/A interneuron (left bottom; interneuron is indicated with a white arrow; images acquired with 25x lens, NA 0.95) and a CA1 O/A VIP-LRP labeled with RetroBeads (right top) and expressing GCaMP6f (right middle, bottom). (G) Histograms of Ca2+-transient peak (%ΔF/F, top) and ratio of ΔF/F peak amplitude during immobility to that during locomotion (bottom) obtained from VIP-LRPs labeled with RetroBeads (n = 5 cells/2 imaging sessions of 5 min each, 25–30 locomotion/immobility periods analyzed per cell). Note that Ca2+-signals recorded during immobility (n = 81 periods) were on average larger in amplitude than those recorded during locomotion (n = 63 periods) state at p<0.01, one-way ANOVA followed by Tukey’s test.

During the experiment, habituated mice showed spontaneous alternations in their behaviour between locomotion (running speed median and interquartile range: 8.8 and 5–30 cm/s), immobility and flickering (Villette et al., 2017), a transitional state associated with brief random movements (Figure 5A, n = 6 mice). As flickering periods were very short (<1 sec) and variable in occurrence, they were excluded from further analysis. Among 54 CA1 VIP+ interneurons imaged, 28 VIP+ cells were located within O/A and analyzed in details during 409 locomotion and 356 immobility periods (10–15 periods/cell pooled from 2 independent imaging sessions of 5 min each/3 mice; Table S4). Consistent with previous observations in neocortical circuits (Fu et al., 2014), as a population, the majority of VIP+ O/A cells were more active during animal locomotion (Table S4). By applying Otsu’s method (Otsu, 1979) to somatic Ca2+-activity, we could segregate these cells onto two distinct sub-types (Figure 5D): type I VIP cells (n = 8), on average, exhibited higher somatic Ca2+-signals during immobility than during locomotion (locomotion: 30.0 ± 13.2% ΔF/F vs immobility: 100.4 ± 28.0% ΔF/F; p < 0.05; Mann-Whitney test; Figure 5D), while type II cells (n = 20) were more active during locomotion than during quiet states (locomotion: 94.9 ± 10.1%ΔF/F vs immobility: 31.7 ± 4.0%ΔF/F; p < 0.001; Mann-Whitney test; Figure 5D). Post hoc immunohistochemical analysis of recorded neurons (6 out of 28 VIP+ O/A interneurons recorded in vivo were found after brain re-sectioning and processed for markers; Figure 5E) revealed that type I cells which were more active during quiet states express M2R but not CR (2 cells out of 2 tested) and, therefore, correspond to VIP-LRP neurons (Figure 5E, 6D). The type II VIP cells tested were negative for both M2R and CR (4 cells out of 4 tested; Figure 6E) and, therefore, could correspond to VIP-BCs or other VIP+ interneurons. To further validate these data, we performed Ca2+ imaging of CA1 O/A VIP+ interneurons that were retrogradely labelled with red RetroBeads injected in SUB in addition to Cre-driven GCaMP6f expression (Figure 5F; n = 5 cells/2 mice). Local delivery of a small volume of RetroBeads in SUB (25 nL; Figure 5F, left top) resulted in labeling of CA1 PCs as well as of a few O/A interneurons (Figure 5F, left bottom), including VIP+ cells, which thus corresponded to VIP-LRPs (5 out of total 11 interneurons labelled with RetroBeads in 2 mice; Figure 5F, right). We confirmed that bead-labelled interneurons exhibit normal physiological properties using patch-clamp current clamp recordings in vitro (Figure S4). In vivo, Ca2+ transients detected in retrogradely labelled VIP-LRPs had larger peak amplitude during immobility than during locomotion periods (immobility: 55.3 ± 6.7% ΔF/F, n = 81 periods/ 5 cells; locomotion: 21.0 ± 3.2% ΔF/F, n = 63 periods/ 5 cells; p<0.01, one-way ANOVA followed by Tukey’s test; Figure 5G, S4C–S4D). Taken together, these data indicate that VIP-LRPs are more active at rest than during locomotion. As VIP-LRP cells showed different levels of somatic activity during behavioural states, we next examined their recruitment during network oscillations (Malvache et al., 2016; Villette et al., 2017). The locomotion periods were associated with prominent theta oscillations (7.1 ± 0.3 Hz; n = 6 mice; Figure 6A–6C), while high-frequency ripples were observed during the animal quiet state (144.5 ± 2.6 Hz; n = 6 mice; Figure 6A–6C). For VIP-LRP population (type I VIP+ cells), we combined the cells segregated based on their behaviour activity pattern (n = 6 cells with LFP recorded; Figure 5D) with those that were labelled retrogradely (n = 5; Figure 5F, 5G). In these cells, the peak somatic Ca2+ signal decreased significantly, from 80.80 ± 9.03% ΔF/F during quiet states to 38.61 ± 5.15% ΔF/F during theta-run epochs (p < 0.001; Mann-Whitney test; 154 stationary and 116 theta-run periods, n = 11 cells; Figure 6B), indicating that VIP-LRPs decrease their activity during theta oscillations. In contrast, the M2R-/CR-type II VIP+ cells increased their activity from 31.8 ± 3.3% ΔF/F during quiet state to 96.8 ± 6.1% ΔF/F during theta-run episodes (p < 0.001; Mann-Whitney test; 162 stationary and 190 theta-run periods, n = 14 cells; Figure 6C), pointing to the on-going recruitment of these cells during theta. As the quiet state in awake rodents is associated with recurrent ripple oscillations (Katona et al., 2014; Lapray et al., 2012; Varga et al 2012) and these events may co-occur in the two hippocampi (Buzsaki et al., 1992; 2003; Chrobak and Buzsaki, 1996; Malvache et al., 2016, but see Villalobos et al., 2017 for rats during sleep), we investigated the potential recruitment of VIP+ neurons during ripple episodes. The VIP-LRPs showed no change in somatic activity in relation to ripple episodes (57.2 ± 34.1% ΔF/F before vs 52.7 ± 29.3% ΔF/F after ripple episode, n = 56 episodes/5 cells; p > 0.05; Mann-Whitney test; Figure 6B). Similar data was obtained for the type II M2R-/CR-VIP+ cells (22.2 ± 4.5% ΔF/F before vs 23.5 ± 6.2% ΔF/F after ripple episode, n = 13; p > 0.05; Mann-Whitney test; Figure 6C), although strong ripple coupling was observed in some M2R-/CR- VIP-expressing cells located within PYR (Figure S5). Collectively, these data reveal the preferential recruitment of VIP-LRPs during quiet wakefulness and the suppression in their activity during theta-run epochs, pointing also to functional segregation of VIP interneuron sub-types during different behavioural and network states.

**Fig. 6.**
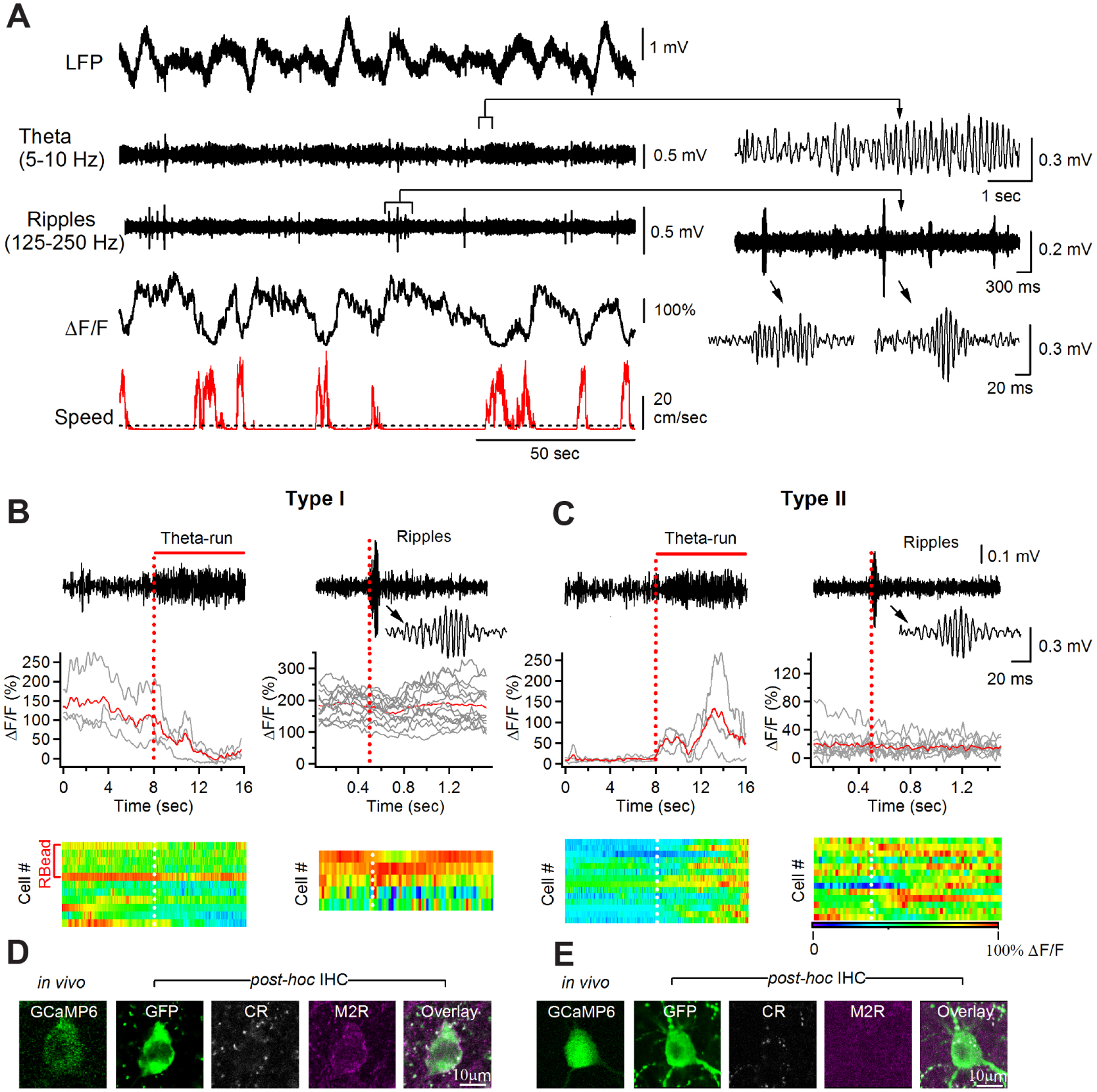
Network state-dependent recruitment of VIP-OA interneurons in awake mice. **See also Figure S4**. (A) Representative traces of simultaneous LFP (raw trace and filtered for theta and ripples) and Ca2+-transient (ΔF/F) recordings from a putative VIP-LRP cell identified post hoc as M2R-positive (D). Red trace illustrates the animal locomotion speed (dotted line indicates the threshold for the locomotion state at 2 cm/sec). (B) Individual traces from the event-triggered Ca2+-trace segmentation and corresponding average (red trace) generated by the theta-run epochs (left) and ripple episodes (right; with inset showing an expanded view of the ripple event.) from the cell illustrated in (A) with heat-maps showing the group data for all VIP-LRPs (n = xx events/6 cells for theta-run; n = xx events/5 cells for ripples). Decrease in somatic Ca2+-signals was significant during theta-run epochs for a group of cells (n = 6) at p< 0.001; Mann-Whitney test. (C) Representative traces from the event-triggered Ca2+-trace segmentation with corresponding average (red trace) generated by the theta-run epochs (left) and ripples (right) with heat-maps showing the group data for type II M2R-/CR-VIP-expressing cells (n = 14 cells for theta-run; n = 13 cells for ripples). Increase in somatic Ca2+-signals during theta-run epochs was significant for a group of cells (n = 14) at p < 0.001; Mann-Whitney test. (D) Post hoc immunohistochemical analysis of the recorded VIP-LRP showing that cells of this sub-type (type I) express M2R. GFP was revealed with Alexa-488, CR with Cy3 and M2R with CF-633 secondary antibodies. (E) Post hoc immunohistochemical analysis of the recorded type II VIP-expressing cells showing that cells of this type do not express M2R or CR.

### Diversity of subiculum-projecting VIP-LRPs

How diverse is the population of SUB-projecting VIP-LRPs? To address this question, we conducted retrograde tracing by (1) injecting a small volume (20 nL) of Cre-dependent hEf1-LS1L-GFP herpes simplex virus (HSV) into the SUB of VIP-Cre;Ai9 mice (Figure 7A, 7B) or (2) using a combinatorial VIP-LRP targeting via injection of retrograde Cav2-Cre into SUB of VIP-flp;Ai65 mice (Figure 7A, 7C). The reporter Ai65D (B6;129S-Gt(ROSA)26Sortm65.1(CAG-tdTomato)Hze/J) mouse line expresses tdTomato under the control of cre and flp. Crossing this reporter line with VIP-flp mice and injecting the Cav2-Cre in the SUB allows selective targeting of VIP+ SUB-projecting neurons. With both strategies, prior calibration experiments were performed to control for the virus spread from SUB to CA1 (Figure 7A; see Methods for details). In addition to a small population of local SUB VIP+ cells (Figure 7B, left), CA1 VIP+ interneurons with somata located within O/A, PYR, RAD or LM were sparsely labelled (VIP-Cre;Ai9 mice + HSV-GFP: 6.7 ± 0.4% of total CA1 VIP+ population, 103/1522 cells from 3 animals, Figure 7A, 7B; VIP-flp;Ai65 + Cav2-Cre: 7.3 ± 0.6% of total CA1 VIP+ population, 100/1364 cells from 3 animals, Figure 7A, 7C). O/A VIP-LRPs made 20% of the total VIP-LRP population (22 out of 103 cells in VIP-Cre;AI9 mice). Consistent with our findings of a low fraction of M2R+ VIP O/A cells in VIP-Cre mice (Figure S3E), some O/A VIP-LRPs labelled with an HSV-GFP in VIP-Cre;Ai9 mice co-expressed M2R (Figure 7D, left; 2 out of 13 O/A VIP-LRPs tested). In addition, those with soma located within PYR, RAD or LM co-expressed CR (10 out of 31 cells tested; Figure 7D, middle) or proenkephalin (Penk, 2 out of 28 cells tested; Figure 7D, right), revealing further molecular diversity within the SUB-projecting VIP-LRP population.

**Fig. 7.**
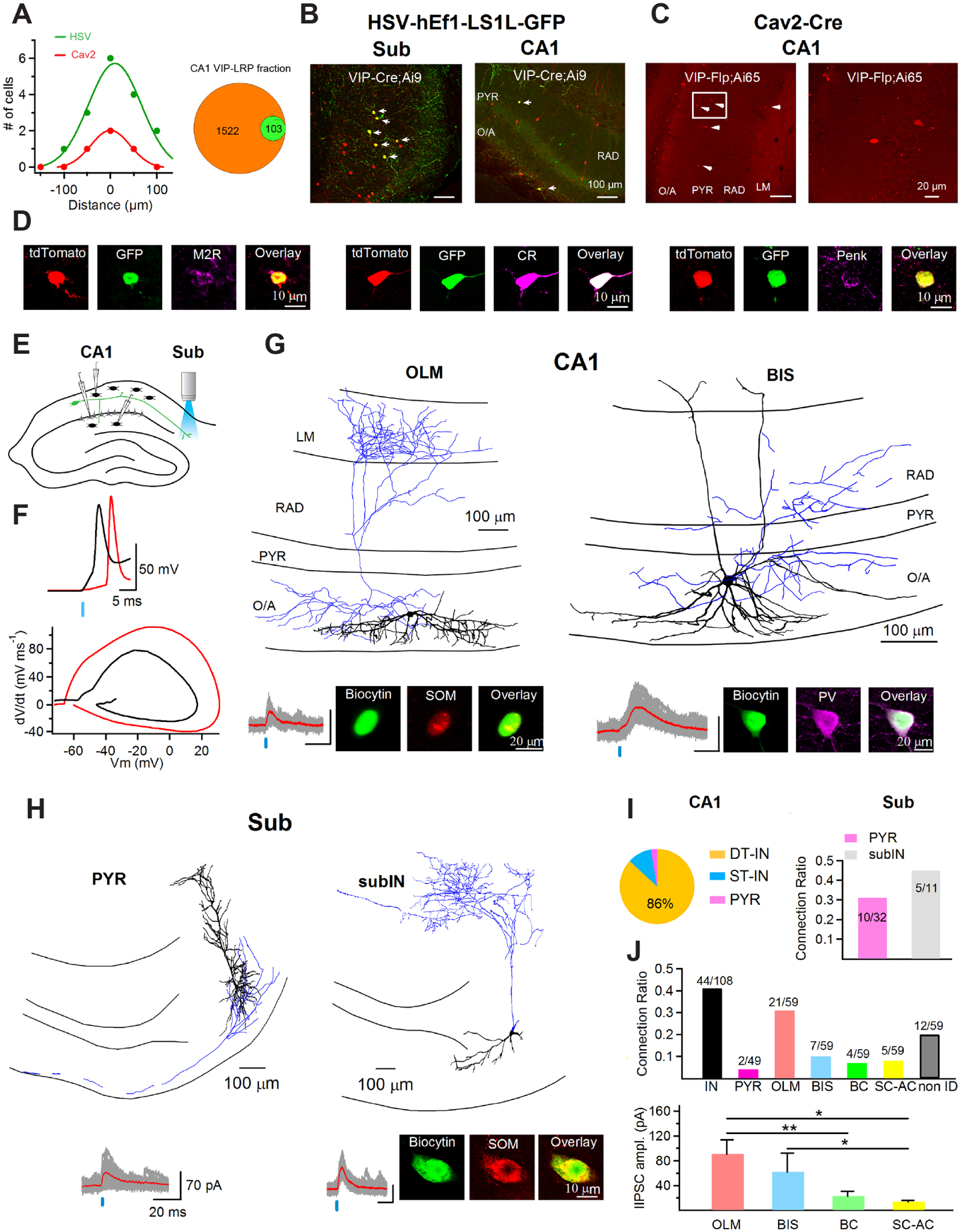
Cellular diversity and connectivity of subiculum-projecting VIP-LRPs. (A) Summary plot (left) for the distribution of infected neuronal cell bodies within SUB for two retrograde viruses (HSV-hEf1-LS1L-GFP and Cav2-Cre) following injection (20 nl total volume) in SUB (the “zero” distance corresponds to the slice taken from the focus of the virus injection; each data point spaced 50 µm apart indicates the total number of labelled cells in the adjacent slice), and a summary pie-chart illustrating the fraction of CA1 VIP-LRPs (green) out of the total VIP+ population (orange) in the CA1 area of VIP-Cre;Ai9 mice injected with a Cre-inducible herpes simplex virus (HSV-hEf1-LS1L-GFP; right). (B) Representative confocal images showing retrograde labelling of subiculum-projecting VIP-LRPs (right) using an HSV-hEf1-LS1L-GFP injection in the subiculum of VIP-Cre;Ai9 mice (left). Red signal – tdTomato-expressing VIP+ neurons; yellow signal – a subpopulation of GFP-VIP+ cells (indicated with white arrowheads) that were labelled with the virus within the subiculum (left) or retrogradely in the CA1 (right). (C) Combinatorial genetic labeling of VIP-LRPs using Cav2-Cre virus injections in the subiculum of VIP-Flp;Ai65 mice confirms the location of VIP-LRPs in different CA1 layers. Left, white arrows point to tdTomato-VIP somata expressing tdTomato under cre and flp control. The area indicated with a white rectangle on the left is shown expanded on the right. (D) Representative confocal images illustrating the markers expressed by VIP-LRPs: M2R (left), CR (center) and Penk (right) in VIP-Cre;Ai9 mice. (E) Schematic illustration of the optogenetic activation of VIP-LRPs through light stimulation in the subiculum in parallel with whole-cell patch-clamp recordings in CA1. (F) The antidromic spike (top, red trace) that was evoked in VIP-LRP in response to light stimulation in subiculum in comparison with somatically evoked spike (top, black trace) in response to CA1 light stimulation; Bottom, summary phase plot for antidromic vs somatic spike comparison. (G-H) Light-evoked IPSCs in response to antidromic activation of VIP-LRPs in different CA1 (G) and subicular (H) targets identified anatomically and neurochemically, including an O-LM cell (G, left) and a BIS cell (G, right) in the CA1 area, as well as a pyramidal cell (H, left) and a SOM-positive interneuron (H, right) in the subiculum. Images at the bottom of each panel show immunoreactivity for SOM or PV. (I) Pie chart (left) illustrating the distribution of VIP-LRP targets in CA1 (DT-IN, dendrite-targeting interneuron; ST-IN, soma-targeting interneuron; PYR, pyramidal cell) and summary bar graphs showing the connection ratio for different postsynaptic targets in subiculum (right; PYR – subicular pyramidal cell, subIN – subicular interneuron). The connection ratio is a ratio between the number of connected pairs of a given type and the total number of attempts. (J) Summary bar graphs illustrating the connection ratio for different types of neurons in the CA1 (top) and the lIPSC amplitude in different CA1 interneuron targets (bottom).

To examine the local connectivity of the entire SUB-projecting VIP+ population in the CA1 area, we next employed a ChR2-assisted circuit mapping approach based on the antidromic activation of VIP-LRP cells through wide-field stimulation of their axons in the SUB of VIP-Cre;Ai32 mice (Figure 7E, 7F). Importantly, we found no evidence for the existence of subiculo-hippocampal VIP+ projecting neurons that could be activated by light-stimulation in SUB and contact CA1 interneurons (Figure S6). In addition, no antidromic spikes were evoked in VIP-BCs (n = 3) or IS3 cells (n = 3) by light stimulation in SUB (data not shown), thus validating our photostimulation approach for antidromic activation of hippocampo-subicular VIP-LRP neurons. In total, 59 CA1 interneurons and 49 CA1 PCs were examined as potential VIP-LRP targets. In CA1 O/A, 33 out of 36 interneurons tested were connected and 26 were visualized with biocytin (Figure 7G), including O-LM (n = 21) and BIS (n = 7) cells (Figure 7G, 7J). Moreover, a putative LRP cell, that was negative for SOM and M2R, with a partially myelinated axon traveling outside the hippocampus, received inhibitory input from VIP-LRP neurons (Figure S7A). In CA1 RAD, 12 out of 23 interneurons tested received input from VIP-LRPs, including CCK-expressing Schaffer-collateral-associated cells (n = 5) and BCs (n = 4; Figure S7B, S7C, 7J). The amplitude of light-evoked IPSCs (lIPSCs) was substantially higher in O/A (88. 6 ± 18.3 pA, n = 26) than in RAD interneurons (30.5 ± 7.6 pA, n = 8; P < 0.01, Mann-Whitney test), with the SOM+ O-LM and BIS cells demonstrating the largest amplitude of lIPSCs (Figure 7J). In contrast, out of 49 PCs tested, only 2 cells with soma within O/A received input from VIP-LRPs (Figure S7D, S7E). These data further support the preferential interneuron innervation by VIP-LRP population, revealing the CA1 circuit disinhibition as a local function of SUB-projecting VIP-LRPs. Subicular targets were also examined using a ChR2-assisted mapping strategy but through wide-field photostimulation in CA1 and patch-clamp recordings in subiculum. Out of 43 attempts, 15 subicular neurons were connected to VIP-LRPs (Figure 7H, 7I). Morphological analysis of cells filled with biocytin showed that SUB targets of VIP-LRPs included both PCs (n = 10 out of 32 attempts; Figure 7H, left; 7I) and interneurons (n = 5 out of 11 attempts; one was identified as SOM+, Figure 7H, right; 7I), thus pointing to a shared VIP-LRP input by PCs and interneurons in the distant projection area. Taken together, these data highlight the region-specific target preference of VIP-LRP population and suggest their potential functional role in setting up CA1 disinhibition concurrently with an inhibitory reset in the subiculum.

## Discussion

We discovered a novel population of hippocampal VIP-expressing GABAergic neurons that exhibit specific molecular properties and, in addition to local innervation of CA1, also make long-range projections to the subiculum with region-specific connectivity patterns. These cells are only weakly active during theta oscillations associated with locomotion but maintain high activity level during quiet state. The latter may promote disinhibition of CA1 PCs in parallel with inhibition–disinhibition periods in the subiculum due to a concomitant innervation of both subicular PCs and interneurons. The likely role of VIP-LRP neurons is therefore to synchronize PC ensembles along the hippocampo-subicular axis that may be necessary for memory consolidation during animal quiet state.

We show that VIP-expressing neurons in the mouse CA1 hippocampus form several functionally and molecularly distinct populations, including BCs, local circuit IS cells and LRP neurons. The VIP-LRP cell is a novel circuit element to be included in the CA1 connectome. This cell type is different from IS3 interneurons, which express CR, exhibit distinct electrophysiological parameters and have a similar axon distribution within the CA1 but target local inhibitory neurons (Chamberland et al., 2010; Tyan et al., 2014). Indeed, the major and, perhaps, the most striking feature of VIP-LRPs is their distant projection and innervation of both interneurons and PCs in the distal projection area.

We demonstrate that, locally, VIP-LRP neurons prefer to make synapses with different classes of inhibitory interneurons, either in the O/A or in the RAD. Interneurons that are known to innervate the PC dendrites, including the O-LM, the BIS and the SC-AC cells, were among the targets of VIP-LRP axons. In addition, perisomatic terminating BCs were also innervated. As the activity of VIP-LRPs was strongly decreased during theta-run epochs, these cells are unlikely to modulate CA1 interneuron firing during theta oscillations associated with locomotion. Indeed, in agreement with our observations, most interneurons exhibit a time-locked maximal activity during theta oscillations necessary for the temporal sequence representation in PC firing (Katona et al., 2014; Klausberger et al., 2003; 2004; Lapray et al., 2012). Interestingly, Buzsáki et al. (1983) identified some rare cells in the hilus as ‘anti-theta’ cells, which were later found in the CA1, subiculum and entorhinal cortex, and classified as theta-off cells (Colom and Bland, 1987; Colom et al., 1987; Mizumori et al., 1990). Despite the remarkable network behavior of theta-off cells, their cellular identity and connectivity patterns have remained unknown. We provide evidence that, at least in the CA1 hippocampus, theta-off cells include a population of VIP-LRP GABAergic neurons that mediate local disinhibition. Importantly, the theta-off cells can display tonic firing by inactivation of the MS (Mizumori et al., 1990), pointing to critical MS suppressive influences in their network motif. In particular, the activation of the M2Rs expressed in the somato-dendritic membrane of these cells or at presynaptic excitatory terminals may be responsible for suppression in VIP-LRP activity during theta oscillations (Apostol and Creutzfeldt, 1974; Fukudome et al., 2004; Lawrence et al., 2015). Furthermore, the activation of local or long-range GABAergic projections (Acsady et al., 1996b; Kaifosh et al., 2013; Unal et al., 2015) that likely converge onto VIP-LRP neurons may prevent these cells from firing during theta-run epochs. Our data also indicate that, in response to the theta-modulated input, VIP-LRPs show weak synchrony due to surround inhibition resulting from the preferential propagation of the spike AHP through gap junctions (Vervaeke et al., 2010). Therefore, on the contrary to other cell types (Mann-Metzer and Yarom, 1999; van Welie et al., 2016), electrotonic coupling between VIP-LRPs does not promote their synchrony; at least this is not the case in response to the theta-like input. How the different firing frequency, the state of gap junctions or the number of coupled neurons participating to the network activity (Alvarez et al., 2002; Trenholm et al., 2014; Pernelle et al., 2018) may shape the cell recruitment remains to be determined.

Are VIP-LRPs discovered here similar to other subiculum-projecting hippocampal GABAergic neurons? One candidate is the trilaminar cell identified previously in the rat hippocampus (Sik et al., 1995), which was shown to express M2R in the somato-dendritic membrane, was decorated with mGluR8a-containing terminals and projected to the subiculum (Ferraguti et al., 2005). This cell had a large soma with horizontally running dendrites at the O/A border and an axon innervating the CA1 from O/A through PYR to RAD. However, the distribution of the trilaminar cell axon in the rat was biased toward proximal RAD (70%; Sik et al., 1995), which is not the case for VIP-LRP cells in the mouse. The presence of VIP has not been reported in the trilaminar cell, and, in contrast to VIP-LRPs, this cell shows complex spike bursts during theta oscillations and strong discharges during ripples (Ferraguti et al., 2005). The other populations of subiculum-projecting GABAergic neurons were described in the outer molecular layer of the dentate gyrus (Ceranik et al., 1997) and in the CA1 (Fuentealba et al., 2008; Jinno et al., 2007). The latter express COUP-TFII alone or in combination with enkephalin or calretinin, but do not express VIP. Moreover, in contrast to VIP-LRPs, these subiculum-projecting GABAergic neurons are strongly modulated during theta oscillations. In conclusion, our data identify the VIP-LRP neuron as a novel circuit element, which, through its region- and target-specific GABAergic interactions, controls the information flow along the hippocampo-subicular axis. As activation of VIP-LRPs occurred preferentially during animal quiet state, this cell type constitutes a good candidate for hippocampo-subicular mnemonic processing associated with episode recollection and comparison (Deadwyler and Hampson, 2004; Kim et al., 2012). Indeed, the coherency between the two regions increases during quiet network states (Jackson et al., 2011; 2014) and, in addition to other mechanisms, may require the involvement of hippocampo-subicular VIP-LRP GABAergic neurons.

## METHODS

### Mouse lines

Nine mouse lines were used in this study: the previously characterized VIP/enhanced green fluorescent protein (VIP-eGFP; Tyan et al., 2014) mice [BAC line with multiple gene copies; MMRRC strain 31009, STOCK Tg(Vip-EGFP) 37Gsat, University of California, Davis, CA]; the previously described VIP-Cre mice (stock 010908, The Jackson Laboratory; Taniguchi et al., 2011; see Supplemental References); the previously characterized VIP-Cre;Ai32 mice (David and Topolnik, 2017), which were obtained by breeding the VIP-Cre with the Ai32 line (B6;129S-Gt(ROSA)26Sortm32(CAG-COP4*H134R/EYFP)Hze/J; stock 012569, The Jackson Lab); Vip-Cre;Ai9 mice obtained by breeding the VIP-Cre mice with the reporter line Ai9-(RCL-tdTomato)line(B6.Cg-Gt(ROSA)26Sortm9(CAG-tdTomato)Hze/J, stock 007909, The Jackson Laboratory) and Vip-Flp;Ai65 mice obtained by breeding the VIP-FlpO mice (kindly provided by Dr. Ed Callaway under agreement with Dr. Josh Huang, CSHL) with a combinatorial reporter Ai65D line (B6;129S-Gt(ROSA)26Sortm65.1(CAG-tdTomato)Hze/J, stock 021875, The Jackson Laboratory). In VIP-eGFP mice, virtually all interneurons that were immunoreactive for VIP endogenously were confirmed to express eGFP (Figure S2G, S2I; see also Tyan et al., 2014). In VIP-Cre;Ai9 mice hippocampus (Figure S3), we also confirmed the presence of molecular cell type markers found in VIP-eGFP mouse interneurons, albeit at a different proportion. Mice had access to food and water ad libitum and were housed in groups of two to four. Mice undergoing surgery were housed separately (1/cage). All experiments were conducted in accordance with the Animal Protection Committee of Université Laval and the Canadian Council on Animal Care.

### Viral constructs

The pAAV-Ef1a-DIO-hChR2(H134R)-EYFP-WPRE-pA virus was acquired from the University of North Carolina (UNC) at Chapel Hill Vector Core. The AAV1.Syn.Flex.GCaMP6f.WPRE.SV40 was acquired from the University of Pennsylvania Vector Core. The hEf1-LS1L-GFP HSV vector was provided by Dr. Rachael Neve at the MIT Viral Gene Transfer Core and packaged at the University of Massachusetts Medical School Gene Therapy Center and Vector Core. The Cav2-Cre virus was acquired from the Plateforme de Vectorologie de Montpelier (PVM) at Bio-Campus Montpelier.

### Slice preparation and patch-clamp recordings

Transverse hippocampal slices (thickness, 300 µm) were prepared from VIP-eGFP or VIP-Cre;Ai32 mice of either sex as described previously (Chamberland et al., 2010; Tyan et al., 2014). Briefly, animals (P15-30) were anaesthetized deeply with isoflurane or ketamine-xylazine (ketamine: 100 mg/kg, xylazine: 10 mg/kg) and decapitated. The brain was dissected carefully and transferred rapidly into an ice-cold (0 to 4°C) solution containing the following (in mM): 250 sucrose, 2 KCl, 1.25 NaH2PO4, 26 NaHCO3, 7 MgSO4, 0.5 CaCl2, and 10 glucose oxygenated continuously with 95% O2 and 5% CO2, pH 7.4, 330–340 mOsm/L. Transverse hippocampal slices (thickness, 300 µm) were cut using a vibratome (VT1000S; Leica Microsystems or Microm; Fisher Scientific), transferred to a heated (37.5°C) oxygenated recovery solution containing the following (in mM): 124 NaCl, 2.5 KCl, 1.25 NaH2PO4, 26 NaHCO3, 3 MgSO4, 1 CaCl2, and 10 glucose; pH 7.4; 300 mOsm/L and allowed to recover for 1 h. Subsequently, they were kept at room temperature until use. During experiments, slices were continuously perfused (2 mL/min) with standard artificial cerebrospinal fluid (ACSF) at physiological temperature (30–33ºC) containing the following (in mM): 124 NaCl, 2.5 KCl, 1.25 NaH2PO4, 26 NaHCO3, 2 MgSO4, 2CaCl2, and 10 glucose, pH 7.4 saturated with 95% O2 and 5% CO2. VIP-positive O/A interneurons were visually identified as GFP-expressing cells upon illumination with blue light (filter set: 450–490 nm). Two-photon images of GFP-expressing interneurons in acute slices were obtained using a two-photon microscope (TCS SP5; Leica Microsystems) based on a Ti-Sapphire laser tuned to 900 nm. Images were acquired with a 25x water-immersion objective (NA 0.95). Whole-cell patch-clamp recordings were obtained from single cells or pairs of neurons in voltage-or current-clamp mode. Recording pipettes (3.5–6 MΩ) were filled with a Cs-based solution for voltage-clamp recordings (in mM): 130 CsMeSO4, 2CsCl, 10 diNa-phosphocreatine, 10 HEPES, 4 ATP-Tris, 0.4 GTP-Tris, 0.3% biocytin, 2 QX-314, pH 7.2–7.3, 280–290 mOsm/L.; or a K+-based intracellular solution for current-clamp recordings (in mM): 130 KMeSO4, 2 MgCl2, 10 diNa-phosphocreatine, 10 HEPES, 4 ATP-Tris, 0.4 GTP-Tris and 0.3% biocytin (Sigma), pH 7.2–7.3, 280–290 mOsm/L. Data acquisition (filtered at 2-3 kHz and digitized at 10kHz; Digidata 1440, Molecular Devices, CA, USA) was performed using the Multiclamp 700B amplifier and the Clampex 10.5 software (Molecular Devices). Active membrane properties were recorded in current-clamp mode by subjecting cells to multiple current step injections of varying amplitudes (−240 to +280 pA).

To assess synaptic connectivity between VIP-LRPs and O/A interneurons, two neurons were recorded simultaneously, with the presynaptic interneuron (VIP-LRP) kept in current-clamp mode at –60 mV and the postsynaptic cell (O/A interneuron) held in voltage-clamp mode at 0 mV. The junction potential was not corrected. Action potentials (APs) were evoked in the presynaptic interneuron via 2 brief somatic current injections (2 ms, 1–1.5 nA) at 20 Hz. In case of synaptic connection, this protocol evoked short-latency (<5 ms) unitary IPSCs (uIPSCs) in the postsynaptic cell. Although individual uIPSCs were small (10 pA) and close to the noise level in our experiments (5–9 pA), we could detect them based on a constant latency (Figure 2C, 2F, 2G). The pipette capacitance and series resistance (in voltage-clamp configuration) were compensated and bridge balance (in current-clamp configuration) was adjusted. The series resistance (Rser) before compensation was 15–20 MΩ and was monitored continuously by applying a –5 mV step at the end of every sweep. Recordings with changes in Rser > 15% were removed from the analysis. To detect changes in uIPSCs amplitude during different frequencies of firing of VIP-LRPs, APs were generated in VIP-LRPs at 10 Hz, 50 Hz, and 100 Hz. To examine electrical coupling between VIP-LRPs, two neurons were recorded simultaneously as mentioned above in the presence of synaptic blockers: gabazine (1 µM), NBQX (10 µM) and AP5 (100 µM). The coupling coefficient (CC12) was calculated as the ratio of voltage responses of the receiving cell (here, cell 2) to the stimulated cell (here, cell 1) with a hyperpolarizing current step (−140 pA, 1000 ms) applied to the cell 1. Pairs were considered to be electrically coupled if their coupling coefficient was higher than 0.01 (Parker et al., 2009; see Supplemental References). Gap junctions were tested with a selective connexin-36 gap-junction blocker mefloquine [100 µM, M2319, Sigma; (Cruikshank et al., 2004)] or a broad-spectrum gap-junction blocker carbenoxolone (100 µM, C4790, Sigma). A sinusoidal excitatory input modulated at theta-frequency (5-Hz) was applied to the electrically coupled pair in 3 different conditions: to the cell 1 only with both cells held at resting membrane potential (Figure 3F), to the cell 1 only when cell 2 was depolarized to allow for spontaneous firing (Figure 3G), and to both cells at more depolarized membrane potential (Figure 3H).

### Two-photon laser scanning photostimulation by glutamate uncaging

Two-photon glutamate uncaging experiments were performed as described previously (Chamberland et al., 2010). Briefly, acute hippocampal slices (300 µm) were obtained from VIP-eGFP mice (P15–25) and perfused during experiment with ACSF containing high Ca2+ (4mM), high Mg2+ (4mM) and DL-AP5 (50µM) to reduce the spontaneous synaptic activity and the confounding polysynaptic effects (Shepherd et al., 2003; Brill and Huguenard, 2009; Supplemental References). To avoid non-specific effects of 4-methoxy-7-nitroindolinyl (MNI)-caged glutamate (Glu) on inhibitory synaptic transmission (Fino et al., 2009; Supplemental References), the MNI-Glu (5mM; Tocris) was applied locally by fast micropressure pulses (5psi, 5ms) via a glass pipette with a tip diameter of 2–3µm connected to a pressure application system (PicoSpritzer II; Parker Instrumentation, Fairfield, NJ, USA) and positioned 10µm above the putative VIP-LRP. The putative VIP-LRPs were selected for photoactivation based on their soma location in the CA1 O/A and expression of eGFP. They were visualized for puff-pipette positioning and two-photon somatic glutamate uncaging with a two-photon Dodt infrared scanning gradient contrast technique (Dodt-IRSGC; Chamberland et al., 2010) using a two-photon laser scanning system (Leica TCS SP5 microscope with a 40×, 0.8 NA water-immersion objective; Leica Microsystems) based on a Ti-Sapphire laser tuned to 730nm (laser power measured under the objective, 5–10mW). Focal release of glutamate was accomplished by illuminating the somatic region for 180ms (laser power, 25–30mW) immediately after puff application of the caged compound. These settings were reliable in evoking a single spike in VIP+ O/A interneurons. To prevent photodamage, the stimulations were repeated once every 30s and the laser power did not exceed 40mW (measured under the objective). Control experiments included the application of MNI-Glu without subsequent uncaging, and uncaging without prior application of MNI-Glu (Chamberland et al., 2010).

### ChR2-based mapping of VIP-LRP targets

Optogenetic activation of VIP-LRPs was achieved through wide-field low-intensity stimulation with blue light (filter set: 450–590 nm; average power at the sample, 1.30 mW; pulse duration 2.5 or 5 ms, which was corresponded to the minimal duration able to evoke the response) using a 40x water-immersion objective (NA 0.8), which was applied to an area of 0.2 mm2 within the SUB 1.0–1.2 mm away from the CA1 border to generate antidromic spikes in VIP-LRPs while avoiding the activation of VIP-BCs and IS3 cells in CA1. The generation of antidromic spikes was confirmed using current-clamp recordings from VIP-positive O/A neurons in VIP-Cre;Ai32 mice and photostimulation in SUB. The antidromic spike generated in this case differed from the somatic one evoked by light illumination in the CA1 area (Figure 7F). The opposite experimental paradigm, with patch-clamp recordings in the SUB and photostimulation in the CA1, was applied to investigate the targets of VIP-LRPs in subiculum. The light-evoked IPSCs (IPSCLs) were recorded at 0 mV. Using a low-light stimulation paradigm allowed for spatially localized excitation, which generated both successful lIPSCs and failures (Figure 7G, 7H; S7A, S7C, S7D).

### *In vitro* patch-clamp data analysis

Analysis of electrophysiological recordings was performed using Clampfit 10.6 (Molecular Devices) and Igor Pro 6.2 (WaveMetrics). For the analysis of the AP properties, the first AP appearing at current pulse of +40–60 pA within a 50-ms time window was analysed. The AP amplitude was measured from the threshold to the peak. The AP latency was measured from the beginning of the current pulse to the AP threshold level. The AP half-width was measured at the voltage level of the half of AP amplitude. The fast afterhyperpolarization amplitude was measured from the AP threshold. Ih-associated voltage rectification was determined as the amplitude of the membrane potential sag from the peak hyperpolarized level to the stable level when hyperpolarized to –100 mV.

To analyze the properties of uIPSCs, 100 sweeps were acquired. Sweeps with spontaneous activity occurring right before or during uIPSCs were removed. The failures were identified from individual sweeps as traces that did not contain any time-dependent signal after the end of the presynaptic AP. The failure rate was calculated as the number of failures divided by the total number of traces. After this step, all sweeps containing failures were removed and successful uIPSCs were averaged to obtain uIPSC potency for further analysis. The uIPSC latency was determined as the time interval between the peak of a presynaptic AP and the onset of the uIPSC in the postsynaptic cell. The rise time of uIPSC was taken at 20–80% and a monoexponential decay time constant was determined. We did not attempt to calculate the uIPSC synaptic conductance since the GABA reversal potential can be different at different targets and was not examined in this study. The paired-pulse ratio was determined as the ratio between the mean peak amplitude of the second response and the mean peak amplitude of the first response, which were obtained 50 ms apart, including failures. During repetitive stimulation (10–100 Hz; Figure 2C), the peak amplitudes of individual uIPSCs were extrapolated from the baseline by fitting the decay of the preceding uIPSC at the average trace. The connection ratio for specific postsynaptic targets (OLM, BIS and BC; Figure 2I) was determined as a ratio between the number of connected cells of a specific type to the total number of recording attempts (n = 118).

For ChR2-based mapping analysis, the potency of the light-evoked IPSCs (lIPSCs) was determined as the average lIPSC obtained after removal of all sweeps containing failures. The connection ratio for each specific target was determined as described above for paired recordings.

For electrical coupling analysis, cross-correlation functions in Clampfit were used to explore synchrony in voltage fluctuations (Figure 3F) or firing (Figure 3G, 3H) between electrically coupled VIP-LRPs. For spike synchrony, cross-correlation analysis was performed on high-pass (at 125 Hz) filtered voltage traces following the spike detection algorithm to correlate spike start times between the two connected cells.

### Cell reconstruction and immunohistochemistry

For post hoc reconstruction, neurons were filled with biocytin (Sigma) during whole-cell recordings. Slices with recorded cells were fixed overnight with 4% paraformaldehyde (PFA) at 4 °C. To reveal biocytin, the slices were permeabilized with 0.3% Triton X-100 and incubated at 4 °C with streptavidin-conjugated Alexa-488 or Alexa-546 (1:1000) in TBS. For combined morphological and immunohistochemical analysis of recorded cells, the duration of whole-cell recordings was reduced to 10 min, and the concentration of biocytin was increased to 0.5% for reliable axonal labelling. This procedure was not required for the analysis of the expression of the membrane-bound proteins (e.g., M2R, mGluR1). All immunohistochemical tests were performed on free-floating sections (40 or 70 µm thick) obtained with Leica VT1000S or PELCO EasySlicer vibratomes from 3–4 mice (20 sections/animal) per condition. VIP-eGFP, VIP-Cre (injected with GCaMP6f) or VIP-Cre;Ai9 mice were perfused with 4% PFA and the brains were sectioned. Sections were permeabilized with 0.25% Triton X-100 in PBS and incubated overnight at 4 °C with primary antibodies followed by the secondary antibodies. The list of primary and secondary antibodies used is provided in the Supplemental Information (Table S2). For proenkephalin immunoreaction, biotinylation was performed to enhance the labeling specificity. Briefly, following overnight incubation of sections with rabbit proenkephalin primary antibody, biotinylated anti-rabbit antibody was applied for 24 h followed by streptavidin-conjugated AlexaFluor (1:1000; Table S2). For controlling method specificity, the primary antibodies were omitted and sections incubated in the full mixture of secondary antibodies. Under such conditions no selective cell labeling was detected. Confocal images were acquired sequentially using a Leica TCS SP5 imaging system coupled with a 488-nm argon, a 543-nm HeNe and a 633-nm HeNe lasers. Z-stacks of biocytin-filled cells were acquired with a 1-µm step and merged for detailed reconstruction in Neurolucida 8.26.2. The axon length was measured without shrinkage correction. For Figure 1F, an LSM710 confocal microscope (Axio Imager.Z1, Carl Zeiss) with ZEN 2008 software v5.0 (Zeiss) was used to acquire multi-channel fluorescence images sequentially with a DIC M27 Plan-Apochromat 63× (NA 1.4) objective, as described (Viney et al., 2013, see Supplemental References). The cells were considered immunopositive when the corresponding fluorescence intensity was at least twice of that of the background. For representation only, the overall brightness and contrast of images were adjusted manually. Portions of images were not modified separately in any way. As the antibody to detect immunoreactivity for mGluR8 is sensitive to fixation conditions, we used sections from one well-reacting mouse.

### Retrograde labeling

VIP-eGFP, VIP-Cre;Ai9 or VIP-flp;Ai65 mice (P30–100) were anesthetised deeply via the intraperitoneal injection of ketamine/xylazine (ketamine: 100 mg/kg, xylazine: 10 mg/kg). After receiving a subcutaneous injection of Buprenorphine SR (0.6 mg/ml, 0.05/30g), animals were placed in a stereotaxic frame (Kopf Instruments) and craniotomy was performed on the right hemisphere. For subicular injections, the following bregma coordinates were used: AP, –3.62 mm; ML, ±2.4 mm; and DV, –1.4 mm or AP, –2.54 mm; ML, ±0.75–0.85 mm; and DV, –1.65–1.85 mm. For injections in hippocampal CA1, the coordinates were: AP, –2.44 mm; ML, ±2.4 mm; and DV, –1.3 mm. The injection pipette, which was attached to a microprocessor-controlled nanoliter injector (Nanoliter 2000; World Precision Instruments), was lowered at a speed of 1 mm/min, and the injection of red IX RetroBeads (Luma Fluor, Inc., a total volume of 25–30 nL) or retrograde viruses (HSV-hEf1-LS1L-GFP, 20 nl; or Cav2-Cre; 50 nl) was performed at a rate of 1 nL/s. For both retrograde viruses tested (HSV and Cav2-Cre), we detected sparse labelling of local subicular reporter-VIP+ interneurons (Figure 7B) in addition to CA1 VIP-LRPs. To restrict virus spread, in prior experiments, we estimated the minimal volume of virus required to infect subicular VIP+ cells within a maximum distance of 200 µm from the injection site. The virus spatial labeling efficacy was estimated from the number of subicular cells infected in consecutive coronal sections (50-µm thickness), with “zero” distance corresponding to the injection site (Figure 7A, left). Ten minutes after the injection, the pipette was slowly withdrawn, the scalp was sutured, and the animals were allowed to recover. Two days (for RetroBeads) or two-three weeks (for HSV and Cav2-Cre) after the injection, the animals were intracardially perfused with 4%-PFA and hippocampal slices were prepared. For all retrograde labelling estimates, only sections from animals with a highly localized subicular injection without spread to the adjoining CA1 area were included in the analysis.

### Electron microscopy

Slices from VIP-eGFP mice containing recorded cells filled with biocytin were re-sectioned to 70 µm, cryoprotected in 20% sucrose solution in 0.1 M PB for at least 3h and freeze-thawed. Sections were washed in 0.1 M PB and incubated with Streptavidin Alexa 488 (1:1000) in TBS for 48 hours at 4°C. After revealing the biocytin in recorded cells, sections were incubated in biotin (1:100, Vector Labs) overnight followed by the avidin/biotin complex (1:100; Vector Labs) in TBS at 4°C for 48 hours. Sections were reacted with a solution of 0.05% diaminobenzidine and 0.002% hydrogen peroxide (HRP reaction) in Tris buffer for 10 min. After washing in 0.1 M PB, sections were treated with 1% osmium tetroxide solution in 0.1 M PB for 1 h, washed in PB and dehydrated in a graded series of alcohol (70, 90, 95, and 100%) followed by propylene oxide. Uranyl acetate (1%) was added to the 70% alcohol for 35 min for contrast enhancement. Dehydrated sections were embedded in Durcupan resin (Fluka) and polymerized at 60°C for 2 days. Target areas were cut out from the resin-embedded 70-µm-thick sections and re-embedded for ultramicrotome sectioning. Serial 60-nm-thick sections were cut and mounted on single-slot, pioloform-coated copper grids. Sections were observed with a Philips CM100 transmission electron microscope and electron micrographs were acquired with a Gatan UltraScan 1000 CCD camera. Synaptic junctions were examined in CA1 O/A and RAD. The postsynaptic target identity was determined using published criteria. Briefly, postsynaptic interneuron dendrites receive type 1 (asymmetrical) synapses on the dendritic shafts and show no spines or low spine density. In contrast, postsynaptic PCs receive type 1 synapses on their spines and type 2 symmetrical synapses on their shafts (Gulyas et al., 1999; Megias et al., 2001; Supplemental References).

### Two-photon imaging in awake mice

Two-photon somatic Ca2+-imaging of VIP interneuron activity was performed in head-restrained awake mice running on the treadmill, which consisted of a shock absorber free rotating wheel with minimized brain motion artifacts. The running wheel was equipped with lateral walls for increased animal contentment and coupled with an optical encoder allowing for acquisition of running speed synchronously with electrophysiological signal (Villette et al., 2017). Male adult VIP-Cre mice (25–35 g body weight; P40–100) were injected stereotaxically with AAV1.Syn.Flex.GCaMP6f.WPRE.SV40 (stock diluted 1:4 in PBS; total injection volume 100 nl) into two sites of the CA1 hippocampus using the following coordinates: AP, –2.54 mm, ML, –2.1 mm, DV, –1.3 mm and AP, –2.0, ML, –1.6, DV, –1.3 mm. At 7–10 days after viral injection, mice were anaesthetized deeply with a ketamine–xylazine mixture (ketamine: 100 mg/kg, xylazine: 10 mg/kg), and fixed in a stereotaxic frame. For hippocampal imaging window, a glass-bottomed cannula was inserted on top of the dorsal hippocampus after the cortex aspiration, and secured with Kwik-Sil at the tissue interface and Super-bond at the skull level (Dombeck et al., 2010, Supplemental References). For Ca2+ imaging from the retrogradely labelled VIP+ O/A interneurons, VIP-Cre mice injected with AAV-GCaMP6f in the CA1 were receiving an injection of red RetroBeads (25 nL) in the subiculum (AP, –2.54 mm, ML, –0.85 mm, DV, –1.65 mm) before hippocampal imaging window preparation. A single tungsten electrode for LFP recordings was implanted in the contralateral CA1 hippocampus and a reference electrode was implanted above the cerebellum (Malvache et al., 2016, Villette et al., 2017). The head plate was oriented medio-laterally at 7–13° using a four-axis micromanipulator (MX10L, Siskiyou) and fixed with several layers of Superbond and dental cement. Mice were allowed to recover for several days with postoperative pain killer treatment for 3 consecutive days (buprenorphine, 0.1 mg kg–1; 48 h). Behavioural habituation involved progressive handling by the experimenter for 5–15 min twice per day for a total of three days, with the animal fixation in the apparatus starting from the third day. During experiment, the LFP signal acquisition was performed simultaneously with the optical encoder signal and imaging trigger at a sampling frequency of 10 kHz using the DigiData1440 (Molecular Devices), AM Systems amplifier and the AxoScope software (v10.5, Molecular Devices). Imaging was performed using a Leica SP5 TCS two-photon microscope equipped with two external photomultiplier tubes (PMTs) for simultaneous detection of green (GCaMP6f) and red (RetroBeads) fluorescence and coupled with a Ti:sapphire femtosecond laser (Chameleon Ultra II, Coherent), which was mode-locked at 900 nm. A long-range water-immersion 25× objective (0.95 NA, 2.5 mm working distance) was used for excitation and light collection to PMTs at 12 bits. Image series were acquired at axial resolutions of 2 µm/pixel and temporal resolutions of 30–48 images/sec. Two 5-min long recording sessions were acquired for each cell. The experiment lasted up to 1 h, after which the mouse was placed back in its home cage. The locomotion wheel between different animals was cleaned with tap water. The image and LFP analyses were performed off-line using Leica LAS, Igor Pro (Wavemetrics, Lake Oswego, USA), Clampfit 10.6 and Statistica (StatSoft). For post hoc immunohistochemical analysis of VIP-OA interneurons recorded in vivo, a 3D reconstruction of the hippocampal window imaged in vivo was performed using sequential confocal acquisition and automatic stitching. Following in vivo experiments, animals were perfused with 4% PFA, the brains were removed, re-sectioned to 70 µm and processed for GFP, M2R and CR. Sequential confocal Z-stacks (120–150 stacks in total/imaging window, 2-µm step, 500–700-µm depth from the alveus surface) were acquired using Nikon AR1 MP+ multiphoton microscope equipped with a 20x objective (NA 1.1), and automatic stitching of individual Z-stacks was applied using NIS Elements AR 4.51.00 software (Nikon Instruments).

### Analysis of two-photon Ca2+ imaging data

For the analysis of spontaneous behaviour, three behavioural phases were identified: locomotion, flickering, and immobility. Locomotion epochs were defined as the periods when the instantaneous speed was higher than 2 cm/s for a minimal distance of 2 cm, thereby pooling together the walking and running periods. The periods with small random movements, when the speed was above 0.25 cm/s but below the locomotion threshold, were defined as flickering. Immobility periods were defined as the times without wheel rotation.

The image analysis was performed off-line using Leica LAS, Igor Pro (Wavemetrics, Lake Oswego, USA) and Statistica (StatSoft). Movies were motion corrected along the x-y plane, no neuropil subtraction was performed (Villette et al., 2017). For extraction of somatic Ca2+-transients, a region of interest was drawn around individual soma to generate the relative fluorescence change (F) versus time trace. The baseline fluorescence level (F0) was determined as the average fluorescence signal derived from three 1-sec time intervals corresponding to the lowest fluorescence level in the absence of Ca2+-transients irrespective of the behaviour state. Somatic Ca2+-transients were expressed as %ΔF/F = (F - F0)/F0 x 100%. Peak Ca2+-signals were determined as averaged signals derived from 315-msec windows around the peak of individual Ca2+-transients over a total period of locomotion or immobility. A total of 7–10 individual Ca2+-transients were analyzed per animal behavioural state with a total of 10–15 states per cell to calculate the average peak Ca2+-transient/state for a given cell. This analysis was performed for two independent imaging sessions (S1 and S2, AVG - average between the two; Figure 5D). Ca2+-transient peak amplitudes recorded during two behavioural states (locomotion and immobility) were tested for normality in their distribution using the Shapiro-Wilcoxon test. Otsu’s method based on the discrimination criterion was applied to somatic Ca2+-activity recorded in VIP O/A neurons during locomotion and immobility to identify two types of cells (Figure 5D). The discrimination criterion between the two groups of cells was 0.86, indicating a good separability. Furthermore, the Mann-Whitney test was used to determine whether obtained groups of neurons exhibit statistically different properties. The results of neuron classification are illustrated as a heat-map with the amplitude of somatic Ca2+-fluctuations during different behavioural states color-coded (Figure 5D). To examine somatic Ca2+-fluctuations in relation to network oscillations, LFP traces were band-pass filtered to obtain theta oscillations (5–10 Hz) or ripples (125–250 Hz). The frequency of theta oscillations during locomotion or ripple events that were detected during quiet state was determined using the power spectrum analysis in Clampfit. The onset of the theta-run epoch, which was always associated with an increase in theta power, was defined by the beginning of the locomotion period based on the animal speed trace acquired simultaneously with LFP (Figure 6B, 6C). Ripple events (9–30 events/cell; minimal event duration: 50 ms; Figure 6B, 6C) were selected semi-automatically at 5 SDs above the signal background using Clampfit event search algorithm with a minimal 50-ms spacing interval between individual events. The frequency at the spectral peak of each selected event was confirmed using power spectrum analysis (Sullivan et al., 2011; Supplemental References). Ca2+-trace segmentation triggered by the event onset (theta-run epoch or ripple) as well as event-triggered averages were conducted in IgorPro.

### Statistics

For statistical analysis, distributions of data were first tested for normality with a Kolmogorov–Smirnov or Shapiro-Wilcoxon test (Figures 1E, 2E, 5D; Tables S1, S3, S4). If data were normally distributed, standard parametric statistics were used: unpaired or paired t tests for comparisons of two groups and one-way or repeated-measures ANOVA for comparisons of multiple groups followed by Tukey, Kruskal–Wallis or Chi2 tests (Figures 1E; 2E; Tables S1, S3, S4). If data were not normally distributed, non-parametric statistics were used: Mann–Whitney or Wilcoxon’s matched pairs test for comparisons of two groups and Kruskal–Wallis test or Dunn’s test for comparisons of multiple groups (Table S1). All statistical analysis was conducted in Sigma Plot 11.0, IgorPro 4.0 or Statistica. P-values < 0.05 were considered significant. Error bars correspond to SEM.

## ACKNOWLEDGEMENTS

We acknowledge the Genie Program and the Janelia Farm Research Campus for GCaMP6f: L.L. Looger, PhD, J. Akerboom, PhD, and D.S. Kim, PhD. We thank Ed Callaway (Salk Institute for Biological Studies) for providing VIP-flpO mice (developed in the laboratory of Josh Huang; CSHL). Special thanks to Sarah Côté and Stéphanie Racine-Dorval for their excellent technical assistance; Marie-Josée Wallman and Martin Parent for their help with tissue preparation for electron microscopy. This work was supported by the Canadian Institutes of Health Research (CIHR Grant MOP-137072, MOP-142447) and the Natural Sciences and Engineering Research Council of Canada (NSERC Grant 342292-2012) to L.T., the British Medical Research Council (MC-UU-12024/4) and the Wellcome Trust (Grant 108726) to P.S. V.V. was supported by the Savoy Foundation Postdoctoral fellowship. O.C. was supported by the NSERC PhD fellowship. S.C. was supported by the FRQS MSc fellowship and the NSERC PhD fellowship for research projects conducted in Topolnik lab. The authors declare no conflict of interest.

## Bibliography

1. Acsády, L., Arabadzisz, D., and Freund, T. F. (1996a). Correlated morphological and neurochemical features identify different subsets of vasoactive intestinal polypeptide-immunoreactive interneurons in rat hippocampus. Neuroscience 73, 299–315.

2. Acsády, L., Görcs, T. J., and Freund, T. F. (1996b). Different populations of vasoactive intestinal polypeptide-immunoreactive interneurons are specialized to control pyramidal cells or interneurons in the hippocampus. Neuroscience 73, 317–334.

3. Alvarez, V. A., Chow, C. C., Van Bockstaele, E. J., and Williams, J. T. (2002). Frequency-dependent synchrony in locus ceruleus: Role of electrotonic coupling. Proceedings of the National Academy of Sciences of the United States of America, 99, 4032–4036.

4. Apostol G, Creutzfeldt OD. (1974). Crosscorrelation between the activity of septal units and hippocampal EEG during arousal. Brain Res. 67, 65–75.

5. Atallah, B. V., and Scanziani, M. (2009). Instantaneous modulation of gamma oscillation frequency by balancing excitation with inhibition. Neuron 62, 566–577.

6. Ayzenshtat, I., Karnani, M. M., Jackson, J., and Yuste, R. (2016). Cortical Control of Spatial Resolution by VIP+Interneurons. J. Neurosci. 36, 11498–11509.

7. Bayraktar, T., Welker, E., Freund, T. F., Zilles, K. and Staiger, J. F. (2000), Neurons immunoreactive for vasoactive intestinal polypeptide in the rat primary somatosensory cortex: Morphology and spatial relationship to barrel-related columns. J. Comp. Neurol. 420, 291–304.

8. Buzsáki, G., Lai-Wo, S. L., and Vanderwolf, C. H. (1983). Cellular bases of hippocampal EEG in the behaving rat. Brain Res. Rev. 6, 139–171.

9. Buzsáki, G., Horvath, Z., Urioste, R., Hetke, J., and Wise, K. (1992). High-frequency network oscillation in the hippocampus. Science 256, 1025–1027.

10. Buzsáki, G., Buhl, D. L., Harris, K. D., Csicsvari, J., Czéh, B., and Morozov, A. (2003). Hippocampal network patterns of activity in the mouse. Neuroscience 116, 201–211.

11. Ceranik, K., Bender, R., Geiger, J. R., Monyer, H., Jonas, P., Frotscher, M., Lübke, J. (1997). A novel type of GABAergic interneuron connecting the input and the output regions of the hippocampus. J Neurosci. 17, 5380–94.

12. Chamberland, S., Salesse, C., Topolnik, D., and Topolnik, L. (2010). Synapse-specific inhibitory control of hippocampal feedback inhibitory circuit. Frontiers in Cell. Neurosci. 4, 130.

13. Chen, X. J., Kovacevic, N., Lobaugh, N. J., Sled, J. G., Henkelman, R. M., and Henderson, J. T. (2006). Neuroanatomical differences between mouse strains as shown by high-resolution 3D MRI. NeuroImage 29, 99–105.

14. Chia, R., Achilli, F., Festing, M. F. W., and Fisher, E. M. C. (2005). The origins and uses of mouse outbred stocks. Nature Genetics, 37, 1181.

15. Chrobak, J., and Buzsaki, G. (1996). High-frequency oscillations in the output networks of the hippocampal-entorhinal axis of the freely behaving rat. Neuroscience 16, 3056–3066.

16. Colom, L. V, Ford, R. D., and Bland, B. H. (1987). Hippocampal formation neurons code the level of activation of the cholinergic septohippocampal pathway. Brain Res. 410, 12–20.

17. Colom, L. V, and Bland, B. H. (1987). State-dependent spike train dynamics of hippocampal formation neurons: evidence for theta-on and theta-off cells. Brain Res. 422, 277–286.

18. Cruikshank, S. J., Hopperstad, M., Younger, M., Connors, B. W., Spray, D. C., and Srinivas, M. (2004). Potent block of Cx36 and Cx50 gap junction channels by mefloquine. Proceedings of the National Academy of Sciences, 101, 12364–12369.

19. Dávid, C., Schleicher, A., Zuschratter, W., and Staiger, J. F. (2007). The innervation of parvalbumin-containing interneurons by VIP-immunopositive interneurons in the primary somatosensory cortex of the adult rat. Eur. J. Neurosci. 25, 2329–40.

20. David, L. S., and Topolnik, L. (2017).Target-specific alterations in the VIP-inhibitory drive to hippocampal GABAergic cells after status epilepticus. Exp. Neurol., 292, 102–112.

21. Deadwyler, S. A., and Hampson, R. E. (2004). Differential but Complementary Mnemonic Functions of the Hippocampus and Subiculum. Neuron 42, 465–476.

22. De Marco García, N. V, Karayannis, T., and Fishell, G. (2011). Neuronal activity is required for the development of specific cortical interneuron subtypes. Nature, 472, 351–355.

23. Ferraguti, F., Klausberger, T., Cobden, P., Baude, A., Roberts, J. D. B., Szucs, P., Kinoshita, A., Shigemoto, R., Somogyi, P., and Dalezios, Y. (2005). Metabotropic Glutamate Receptor 8-Expressing Nerve Terminals Target Subsets of GABAergic Neurons in the Hippocampus. J.Neurosci. 25, 10520–10536.

24. Fu, Y., Tucciarone, J. M., Espinosa, J. S., Sheng, N., Darcy, D. P., Nicoll, R.A, Huang, Z.J., and Stryker, M. P. (2014). A cortical circuit for gain control by behavioral state. Cell 156, 1139–52.

25. Fuentealba, P., Tomioka, R., Dalezios, Y., Marton, L.F., Studer, M, Rockland, K., Klausberger T., Somogyi, P. (2008) Rhythmically active enkephalin-expressing GABAergic cells in the CA1 area of the hippocampus project to the subiculum and preferentially innervate interneurons J. Neurosci. 28, 10017–10022.

26. Fukudome, Y., Ohno-Shosaku, T., Matsui, M., Omori, Y., Fukaya, M., Tsubokawa, H., Taketo, M.M., Watanabe, M., Manabe, T. and Kano, M. (2004). Two distinct classes of muscarinic action on hippocampal inhibitory synapses: M2-mediated direct suppression and M1/M3-mediated indirect suppression through endocannabinoid signalling. Eur. J. Neurosci.19, 2682–2692.

27. Gulyas, A. I., Hajos, N., Katona, I., and Freund, T. F. (2003). Interneurons are the local targets of hippocampal inhibitory cells which project to the medial septum. Eur. J. Neurosci.17, 1861–1872.

28. He, M., Tucciarone, J., Lee, S., Nigro, M. J., Kim, Y., Levine, J. M., Kelly, S.M., Krugikov, I., Wu, P., Chen, Y., et al., (2016). Strategies and tools for combinatorial targeting of GABAergic neurons in mouse cerebral cortex. Neuron 91, 1228–1243.

29. Jackson, J., Goutagny, R., and Williams, S. (2011). Fast and Slow Gamma Rhythms Are Intrinsically and Independently Generated in the Subiculum. J. of Neurosci. 31, 12104–12117.

30. Jackson, J., Amilhon, B., Goutagny, R., Bott, J.-B., Manseau, F., Kortleven, C., Bressler, S.T., and Williams, S. (2014). Reversal of theta rhythm flow through intact hippocampal circuits. Nat. Neurosci. 17, 1362–1370.

31. Jackson, J., Ayzenshtat, I., Karnani, M. M., and Yuste, R. (2016). VIP+ interneurons control neocortical activity across brain states. J. Neurophysiol.115, 3008–3017.

32. Jinno, S., Klausberger, T., Marton, L. F., Dalezios, Y., Roberts, J. D. B., Fuentealba, P., Bushong, E.A., Henze, D., Buzsáki, G. and Somogyi, P. (2007). Neuronal Diversity in GABAergic Long-Range Projections from the Hippocampus. J. Neurosci. 27, 8790–8804.

33. Kaifosh, P., Lovett-Barron, M., Turi, G. F., Reardon, T. R., Losonczy, A. (2013). SeptohippocampalGABAergic signaling across multiple modalities in awake mice. Nat Neurosci. 16, 1182–1184.

34. Karson, M. A., Tang, A., Milner, T. A., and Alger, B. E. (2009). Synaptic cross-talk between perisomatic-targeting interneuron classes expressing cholecystokinin and parvalbumin in hippocampus. J. Neurosci. 29, 4140–4154.

35. Katona, L., Lapray, D., Viney, T. J., Oulhaj, A., Borhegyi, Z., Micklem, B. R., Klausberger, T., and Somogyi, P. (2014). Sleep and movement differentiates actions of two types of somatostatin-expressing GABAergic interneuron in rat hippocampus. Neuron 82, 872–886.

36. Kim, S. M., Ganguli, S., and Frank, L. M. (2012). Spatial information outflow from the hippocampal circuit: distributed spatial coding and phase precession in the subiculum. The Journal of Neuroscience 32, 11539–11558.

37. Klausberger, T., Magill, P. J., Cobden, P. M., and Somogyi, P. (2003). Brain-state- and cell-type-specific firing of hippocampal interneurons in vivo. Nature 421, 844–848.

38. Klausberger, T., Márton, L. F., Baude, A., Roberts, J. D. B., Magill, P. J., and Somogyi, P. (2004). Spike timing of dendrite-targeting bistratified cells during hippocampal network oscillations in vivo. Nature Neurosci. 7, 41–47.

39. Lapray, D., Lasztoczi, B., Lagler, M., Viney, T. J., Katona, L., Valenti, O., Hartwich, K., Borhegyi, Z., Somogyi, P., and Klausberger, T. (2012). Behavior-dependent specialization of identified hippocampal interneurons. Nature Neurosci.15, 1265–1271.

40. Lawrence, J. J., Haario, H., and Stone, E. F. (2015). Presynaptic cholinergic neuromodulation alters the temporal dynamics of short-term depression at parvalbumin-positive basket cell synapses from juvenile CA1 mouse hippocampus. J. Neurophysiol. 113, 2408–2419.

41. Lee, S., Kruglikov, I., Huang, Z. J., Fishell, G., and Rudy, B. (2013). A disinhibitory circuit mediates motor integration in the somatosensory cortex. Nature Neurosci.16, 1662–70.

42. Letzkus, J. J., Wolff, S. B. E., and Lüthi, A. (2015). Dis-inhibition, a circuit mechanism for associative learning and memory. Neuron 88, 264–276.

43. Lovett-Barron, M., Turi, G. F., Kaifosh, P., Lee, P. H., Bolze, F., Sun, X. H., Nicoud, J. F., Zemelman, B. V., Sternson, S. M., Losonczy, A. (2012). Regulation of neuronal inputtransformations by tunable dendritic inhibition. Nature Neurosci.15, 423–30, S1-3.

44. Malvache, A., Reichinnek, S., Villette, V., Haimerl, C., and Cossart, R. (2016). Awake hippocampal reactivations project onto orthogonal neuronal assemblies. Science 353, 1280–1283.

45. Mann-Metzer, P., and Yarom, Y. (1999). Electrotonic Coupling Interacts with Intrinsic Properties to Generate Synchronized Activity in Cerebellar Networks of Inhibitory Interneurons. The Journal of Neuroscience 19, 3298 LP-3306.

46. Mekada, K., Abe, K., Murakami, A., Nakamura, S., Nakata, H., Moriwaki, K., Obata, Y., and Yoshiki, A. (2009). Genetic Differences among C57BL/6 Substrains. Experimental Animals 58, 141–149.

47. Melzer, S., Michael, M., Caputi, A., Eliava, M., Fuchs, E. C., Whittington, M. A., and Monyer, H. (2012). Long-Range–Projecting GABAergic Neurons Modulate Inhibition in Hippocampus and Entorhinal Cortex. Science 335, 1506–1510.

48. Miyashita, T., and Rockland, K. S. (2007). GABAergic projections from the hippocampus to the retrosplenial cortex in the rat. Eur.J.Neurosci. 26, 1193–204.

49. Miyoshi, G., Young, A., Petros, T., Karayannis, T., McKenzie Chang, M., Lavado, A., Iwano, T., Nakajima, M., Taniguchi, H., Huang, Z.J., et al. (2015). Prox1 Regulates the Subtype-Specific Development of Caudal Ganglionic Eminence-Derived GABAergic Cortical Interneurons. J. Neurosci. 35, 12869–12889.

50. Mizumori, SJ, Barnes, CA., and McNaughton, BL. (1990). Behavioral correlates of theta-on and theta-off cells recorded from hippocampal formation of mature young and aged rats. Exp. Brain. Res. 80, 365–373.

51. Otsu, N. (1979). A threshold selection method from gray-level histograms, IEEE transactions on systems, man and cybernetics, Vol SMC-9, No.1.

52. Pernelle, G., Nicola, W., and Clopath, C. (2018). Gap junction plasticity as a mechanism to regulate network-wide oscillations. PLOS Computational Biology 14, e1006025.

53. Pfeffer, C. K., Xue, M., He, M., Huang, Z. J., and Scanziani, M. (2013). Inhibition of inhibition in visual cortex: the logic of connections between molecularly distinct interneurons. Nature Neurosci.16, 1068–76.

54. Pi, H.-J., Hangya, B., Kvitsiani, D., Sanders, J. I., Huang, Z. J., and Kepecs, A. (2013). Cortical interneurons that specialize in disinhibitory control. Nature, 503, 521–524.

55. Porter, J. T., Cauli, B., Staiger, J. F., Lambolez, B., Rossier, J., and Audinat, E. (1998). Properties of bipolar VIPergic interneurons and their excitation by pyramidal neurons in the rat neocortex. Eur. J. Neurosci. 10, 3617–3628.

56. Royer, S., Zemelman, B. V, Losonczy, A., Kim, J., Chance, F., Magee, J. C., and Buzsáki, G. (2012). Control of timing, rate and bursts of hippocampal place cells by dendritic and somatic inhibition. Nature Neurosci.15, 769–775.

57. Sik, A., Penttonen, M., Ylinen, A., and Buzsaki, G. (1995). Hippocampal CA1 interneurons: an in vivo intracellular labeling study. J. Neurosci.15, 6651–6665.

58. Soltesz, I. (2006). Diversity in the neuronal machine: Order and variability in interneuronal microcircuits. Oxford Univ. Press.

59. Somogyi, J., Baude, A., Omori, Y., Shimizu, H., El Mestikawy, S., Fukaya, M., Shigemoto, R., Watanabe, M., and Somogyi, P. (2004). GABAergic basket cells expressing cholecystokinin contain vesicular glutamate transporter type 3 (VGLUT3) in their synaptic terminals in hippocampus and isocortex of the rat. Eur. J. Neurosci. 19, 552–569.

60. Somogyi P., (2010). Hippocampus: Intrinsic Organisation. In: Shepherd G, Grillner S (eds) Handbook of Brain Microcircuits. Oxford University Press, p.148–664.

61. Staiger, J. F., Masanneck, C., Schleicher, A., and Zuschratter, W. (2004). Calbindin-containing interneurons are a target for VIP-immunoreactive synapses in rat primary somatosensory cortex. J. Comp. Neurol. 468, 179–189.

62. Trenholm, S., McLaughlin, A. J., Schwab, D. J., Turner, M. H., Smith, R. G., Rieke, F., and Awatramani, G. B. (2014). Nonlinear dendritic integration of electrical and chemical synaptic inputs drives fine-scale correlations. Nature Neuroscience 17, 1759–1766.

63. Tyan, L., Chamberland, S., Magnin, E., Camiré, O., Francavilla, R., David, L. S., Deisseroth, K., and Topolnik, L. (2014). Dendritic inhibition provided by interneuron-specific cells controls the firing rate and timing of the hippocampal feedback inhibitory circuitry. J. Neurosci.34, 4534–47.

64. Unal, G., Joshi, A., Viney, T. J., Kis, V., Somogyi, P. (2015). Synaptic targets of medial septal projections in the hippocampus and extrahippocampalcortices of the mouse. J. Neurosci. 35, 15812–26.

65. van Welie, I., Roth, A., Ho, S. S. N., Komai, S., and Häusser, M. (2016). Conditional Spike Transmission Mediated by Electrical Coupling Ensures Millisecond Precision-Correlated Activity among Interneurons In Vivo. Neuron 90, 810–823.

66. Varga, C., Golshani, P., and Soltesz, I. (2012). Frequency-invariant temporal ordering of interneuronal discharges during hippocampal oscillations in awake mice. PNAS 109, 2726–2734.

67. Vervaeke, K., Lőrincz, A., Gleeson, P., Farinella, M., Nusser, Z., and Silver, R. A. (2010). Rapid Desynchronization of an Electrically Coupled Interneuron Network with Sparse Excitatory Synaptic Input. Neuron 67, 435–451.

68. Villalobos, C., Maldonado, P. E., Valdés, J. L. (2017) Asynchronous ripple oscillations between left and right hippocampi during slow-wave sleep. PLoS One. 201712(2).

69. Villette, V., Levesque, M., Miled, A., Gosselin, B., and Topolnik, L. (2017). Simple platform for chronic imaging of hippocampal activity during spontaneous behaviour in an awake mouse. Scientific Reports 7, 43388.

